# The correlated state in balanced neuronal networks

**DOI:** 10.1101/372607

**Authors:** Cody Baker, Christopher Ebsch, Ilan Lampl, Robert Rosenbaum

## Abstract

Understanding the magnitude and structure of inter-neuronal correlations and their relationship to synaptic connectivity structure is an important and difficult problem in computational neuroscience. Early studies show that neuronal network models with excitatory-inhibitory balance naturally create very weak spike train correlations, defining the “asynchronous state.” Later work showed that, under some connectivity structures, balanced networks can produce larger correlations between some neuron pairs, even when the average correlation is very small. All of these previous studies assume that the local network receives feedforward synaptic input from a population of uncorrelated spike trains. We show that when spike trains providing feedforward input are correlated, the downstream recurrent network produces much larger correlations. We provide an in-depth analysis of the resulting “correlated state” in balanced networks and show that, unlike the asynchronous state, it produces a tight excitatory-inhibitory balance consistent with in vivo cortical recordings.

## I. INTRODUCTION

Correlations between the spiking activity of cortical neurons have important consequences for neural dynamics and coding [1–3]. A quantitative understanding of how spike train correlations are generated and shaped by the connectivity structure of neural circuits is made difficult by the noisy and nonlinear dynamics of recurrent neuronal network models [4–7]. Linear response and related techniques have been developed to overcome some of these difficulties [7–15], but their application to networks of integrate-and-fire neurons models typically relies on a diffusion approximation that requires an assumption of sparse and/or weak connectivity and an assumption that neurons receive uncorrelated, feedforward Gaussian white noise input. However, cortical circuits are densely connected and receive spatially and temporally correlated synaptic input from outside the local circuit [16–19].

An alternative approach to analyzing correlated variability in recurrent neuronal network models is motivated in part by the widely observed balance between excitatory and inhibitory synaptic inputs in cortex [20–27]. When synaptic weights are scaled like 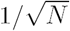 where *N* is the size of a model network, balance between excitation and inhibition arises naturally at large network size, which defines the “balanced state” [28, 29]. A similar scaling of synaptic weights has since been observed in cultured cortical populations [30].

Early work on balanced networks assumed sparse connectivity to produce weak spike train correlations, but it was later shown that keeping connection probabilities 𝒪 (1) naturally produces weak, 𝒪(1*/N)*, spike train correlations, defining the “asynchronous state” [31]. While these extremely weak spike train correlations are consistent with some cortical recordings [32], the magnitude of correlations in cortex can depend on stimulus, cortical area, layer, and behavioral or cognitive state, and can be much larger than predicted by the asynchronous state [6, 33–37]. This raises the question of how larger correlation magnitudes can arise in balanced cortical circuits. Later theoretical work showed that larger correlations can be obtained between some cell pairs in densely connected networks with specially constructed connectivity structure [38–42], offering a potential explanation of the larger correlations often observed in recordings.

All of these previous studies of correlated variability in balanced networks assume that the recurrent network receives feedforward synaptic input from an external population of uncorrelated spike trains, so feedforward input correlations arise solely from overlapping feedforward synaptic projections. In reality, feedforward synaptic input to a cortical population comes from thalamic projections, other cortical areas, or other cortical layers in which spike trains could be correlated.

We extend the theory of densely connected balanced networks to account for correlations between the spike trains of neurons in an external, feedforward input layer. We show that correlations between the feedforward synaptic input to neurons in the recurrent network are 𝒪 (*N)* in this model, but cancel with 𝒪 (*N)* correlations between recurrent synaptic input to produce 𝒪 (1) spike train correlations on average, defining what we refer to as the “correlated state” in densely connected balanced networks. This correlated state offers an alternative explanation for the presence of moderately large spike train correlations in cortical recordings. We derive a simple, closed form approximation for the average covariance between neurons’ spike trains in the correlated state in term of synaptic parameters alone, without requiring the use of linear response theory or any other knowledge of neurons’ transfer functions. We show that the tracking of excitatory synaptic input currents by inhibitory currents is more precise and more similar to in vivo recordings [23] in the correlated state than in the asynchronous state. Our results extend the theory of correlated variability in balanced networks to the biologically realistic assumption that presynaptic neural populations are themselves correlated.

## II. MODEL AND BACKGROUND

We consider recurrent networks of *N* model neurons, *N*_*e*_ of which are excitatory and *N*_*i*_ inhibitory. Neurons are randomly and recurrently interconnected and also receive random feedforward synaptic input from an external population of *N*_*x*_ neurons whose spike trains are homogeneous Poisson processes with rate *r*_*x*_ (Fig. 1A).

**FIG. 1.**
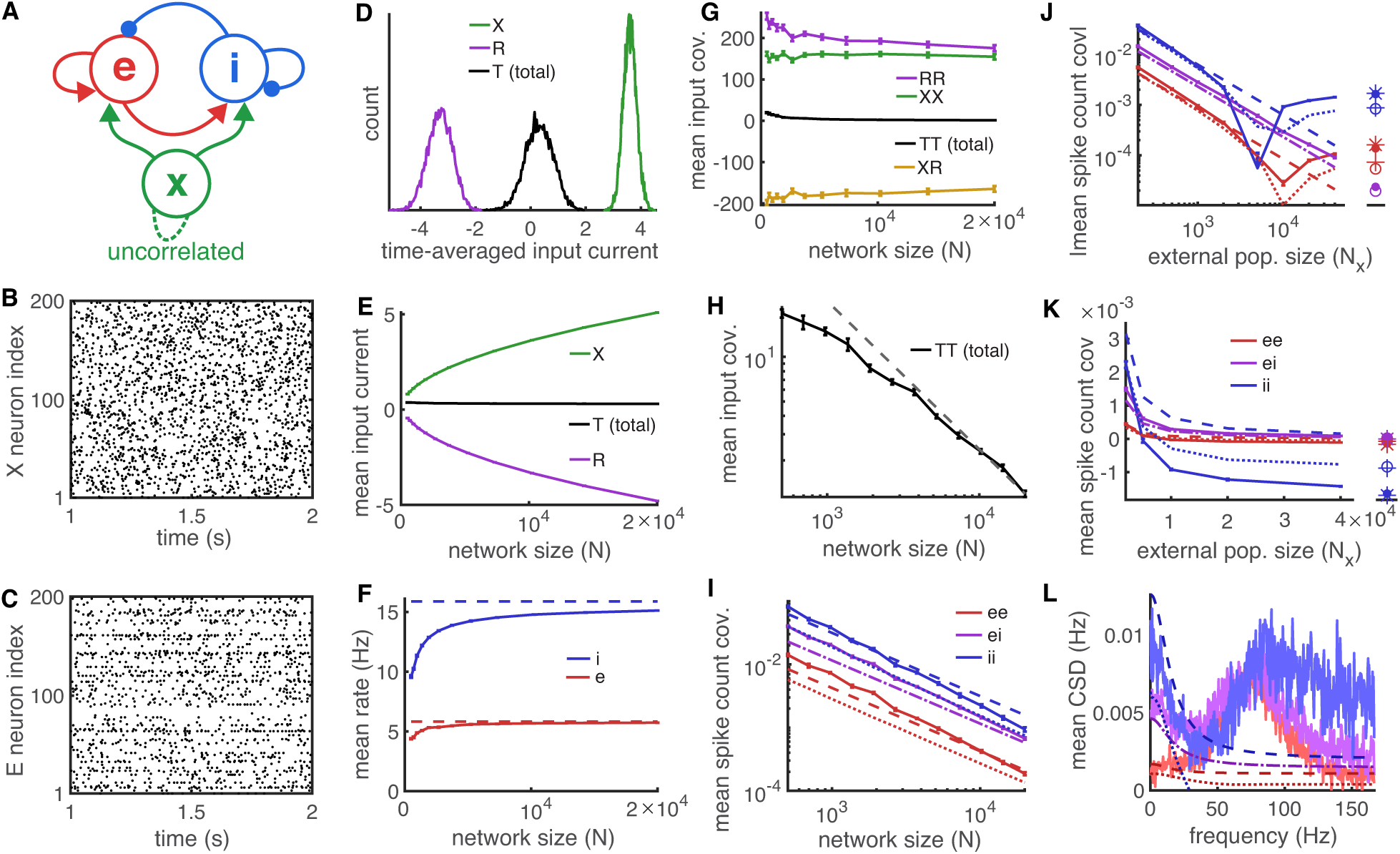
The asynchronous state in densely connected balanced networks. **A)** Network diagram. An external population, *X*, of uncorrelated Poisson processes provides feedforward input to a randomly connected recurrent network of excitatory, *E*, and inhibitory, *I*, neurons. Feedforward input correlations are solely from overlapping projections from *X*. **B,C)** Raster plot of 200 randomly selected neurons from population *X* and *E* respectively in a network with *N* = 10^4^ and *N*_*x*_ = 2000. **D)** Histogram of time-averaged external (*X*, green) recurrent (*R* = *E* + *I*, purple) and total (*T* = *X* + *E* + *I*, black) input to all excitatory neurons in a network with *N* = 10^4^ and *N*_*x*_ = 2000. Currents here and elsewhere are reported in units *C*_*m*_*V/s* where *C*_*m*_ is the arbitrary membrane capacitance. **E)** Mean external (green), recurrent (purple), and total (black) input to excitatory neurons for networks of different sizes, *N*. **F)** Mean excitatory (red) and inhibitory (blue) neuron firing rates for different network sizes. Solid curves are from simulations and dashed curves are from Eq. (5). **G)** Mean covariance between pairs of excitatory neurons’ external inputs (green), recurrent inputs (purple), total inputs (black), and mean covariance between the recurrent input to one excitatory neuron and external input to the other (yellow) for different network sizes. Covariances were computed by integrating the inputs over 250ms windows then computing covariances between the integrals, which is proportional to zero-frequency CSD (see Eq. (3) and surrounding discussion). Integrated currents have units *C*_*m*_*mV*, so their covariances have units 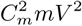. **H)** Zoomed in view of black curve from E on a log-log axis (black) and the function C/N (dashed gray) where C was chosen so that the two curves match at *N* = 2*×*10^4^. **I)** Mean spike count covariance between excitatory neuron spike trains (red), between inhibitory neuron spike trains (blue), and between excitatory-inhibitory pairs of spike trains (purple). In panels I-L, solid curves are from simulations, dashed curves are from the first term in Eq. (9), and dotted curves are from the first two terms. For the dotted curves, the {*S*_*e*_, *S*_*e*_} and {*S*_*i*_, *S*_*i*_} terms in Eq. (9) were estimated empirically from simulations. In all panels, counts were computed over 250ms time windows. **J)** Absolute value of mean spike count covariances as a function of the external population size, *N*_*x*_, when *N* = 10^4^ and where feedforward connection probabilities were scaled to keep *p*_*ex*_*N*_*x*_ = *p*_*ix*_*N*_*x*_ = 200 fixed as *N*_*x*_ was changed. Filled circles are from simulations with uncorrelated external input (representing *N*_*x*_→ *∞*) and open circles are from simulations with deterministic, time-constant external input (external input covariance is zero in both cases). Asterisks and crosses are from evaluating the second term of Eq. (9) using the values of {*S*_*e*_, *S*_*e*_*}* and {*S*_*i*_, *S*_*i*_*}* estimated from the simulations marked with filled circle and crosses respectively. **K)** Same data as J, but plotted on a linear axis without taking absolute value, and zoomed into to larger values of *N*_*x*_. **L)** Mean cross-spectral densities between neurons in the recurrent network using the parameters from panels B-D. Solid curves are from simulations, dashed are from the first term of Eq. (9), and dotted are from the first two terms of Eq. (9) using empirically estimated power spectral densities for {*S*_*b*_, *S*_*b*_*}*(*f)*. Synaptic time constants were *τ*_*e*_ = 8, *τ*_*i*_ = 4, and *τ*_*x*_ = 10ms in all simulations. In all simulations except those in panels J and K, recurrent and feedforward connection probabilities are all *p*_*ab*_ = 0.1 for *a* = *e, i* and *b* = *e, i, x*.

The membrane potential of neuron *j* in population *a* = *e, i* obeys the exponential integrate-and-fire (EIF) dynamics

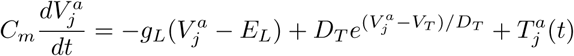

with the added condition that each time 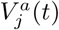 exceeds *V*_*th*_, it is reset to *V*_*re*_ and a spike is recorded. Our results do not depend sensitively on the exact neuron model used or the values of neuron parameters. We additionally set a lower bound on the membrane potential at *V*_*lb*_ = – 100mV. Spike trains are represented as a sum of Dirac delta functions,

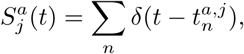

where 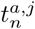 is the *n*th spike time of neuron *j* in population *a* = *e, i, x*. The total synaptic input current to neuron *j* in population *a* = *e, i* is decomposed as

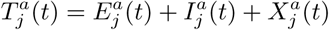

where

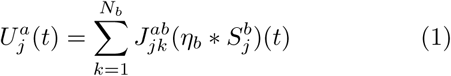

for *U* = *E, I, X* and *b* = *e, i, x* respectively where denotes convolution,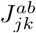 is the synaptic weight from neuron *k* in population *b* to neuron *j* in population *a*, and *η*_*b*_(*t*) is a postsynaptic current (PSC) waveform. Without loss of generality, we assume that ∫*η*_*b*_(*t*) = 1. We use 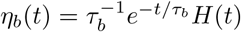 where *H*(*t*) is the Heaviside step function, though our results do not depend sensitively on the precise neuron model or PSC kernel used. For calculations, it is useful to decompose the total synaptic input into its recurrent and external sources,

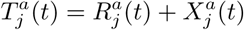

where

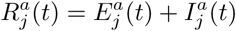

is the recurrent synaptic input from the local circuit.

Local cortical circuits contain a large number of neurons and individual cortical neurons receive synaptic input from thousands of other neurons within their local circuit and from other layers or areas. Densely connected balanced networks have been proposed to model such large and densely interconnected neuronal networks [31, 40]. In such models, one considers the limit of large *N* (with *N*_*x*_, *N*_*e*_ and *N*_*i*_ scaled proportionally) with fixed connection probabilities and where synaptic weights are scaled like 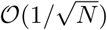 [29, 31]. This scaling naturally captures the balance of mean excitatory and mean inhibitory synaptic input, as well as the tracking of excitation by inhibition, observed in cortical recordings [31]. Recent work in cultured cortical populations shows that similar scaling laws emerge naturally and produce network dynamics consistent with the balanced state [30]. In particular, we consider a random connectivity structure in which

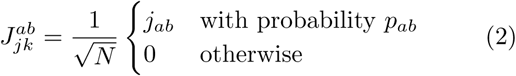

where connections are statistically independent and *j*_*ab*_, *p*_*ab*_ ∼𝒪 (1) for *b* = *e, i, x* and *a* = *e, i*. We furthermore define the proportions

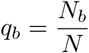

which are assumed 𝒪 (1). For all examples we consider, *q*_*e*_ = 0.8 and *q*_*i*_ = *q*_*x*_ = 0.2.

We next introduce notational conventions for quantifying the statistics of spike trains and synaptic inputs in the network. The mean firing rates of neurons in population *a* = *e, i, x* is defined by *r*_*a*_ for *a* = *e, i, x* and it is useful to define the 2×1 vector, ***r*** = [*r*_*e*_ *r*_*i*_]^*T*^ where .^*T*^ denotes the transpose. The mean is technically interpreted as the expectation over realizations of the network connectivity, but for large *N* it is approximately equal to the sample mean over all neurons the network. Similarly, mean-field synaptic inputs to neurons in populations *a* = *e, i* are defined by

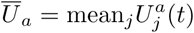

for *U* = *E, I, X, R, T* and, in vector form, 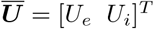

For quantifying correlated variability, we use the crossspectral density (CSD)

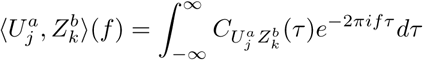

between 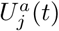 and 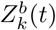 for *U, Z* = *E, I, X, S, R, T* and *a, b* = *e, i, x* where

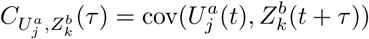

is the cross-covariance function. The argument, *f*, is the frequency at which the CSD is evaluated. The CSD is a convenient measure of correlated variability because it simplifies mathematical calculations due to the fact that ⟨*·,·*⟩, is a Hermitian operator and because most commonly used measures of correlated variability can be written as a function of the CSD. For example, the cross-covariance is the inverse Fourier transform of the CSD. Spike count covariances over large time windows can be written in terms of the CSD by first noting that the spike count is an integral of the spike train [4],

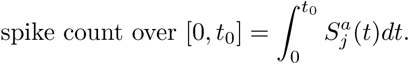

For large *t*_0_, the covariance between two integrals is related to the zero-frequency CSD,

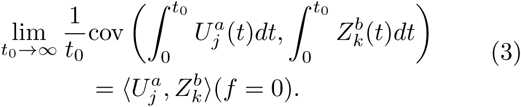

Hence, the spike count covariance over the interval [0, *t*_0_] is given asymptotically by 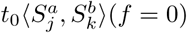 for large *t*_0_. Following this result, we quantify covariability between spike trains and between synaptic currents using the zero-frequency CSD, which we estimate by taking the covariance between integrals as in Eq. (3) using *t*_0_ = 250ms. This provides a simple, easily estimated quantity for quantifying covariance.

Most of our computations are performed at the level of population averages, so we define

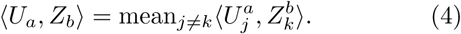

which is a scalar function of frequency, *f*, for each *a, b* = *e, i, x* and *U, Z* = *E, I, X, S, R, T*. It is also convenient to define the 2 × 2 mean-field matrix form,

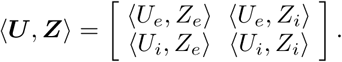

for ***U, Z*** = ***E***, ***I***, ***X***, ***S***, ***R***, ***T***. We also define the recurrent and feedforward mean-field connectivity matrices,

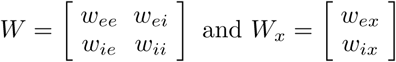

where 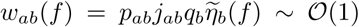 with 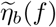 the Fourier transform of *η*_*b*_(*t*). For the exponential kernels we use, 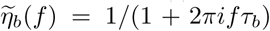. The zero-frequency values,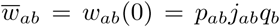, define time-averaged interactions and mean-field connection matrices,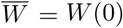 and 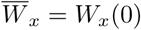.

This choice of notation allows us to perform computations on mean-field spike train and input statistics in a mathematically compact way. We first review the mean-field analysis of firing rates in the balanced state [28, 29, 43–45]. Mean external input is given by 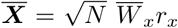 and mean recurrent input by 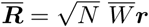 so that mean total synaptic input is given by

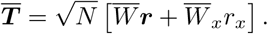

In the balanced state,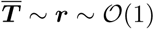, which can only be obtained by a cancellation between externa<u>l a</u>nd r<u>ec</u>urrent synapt<u>ic</u> inputs. This cancellation requires 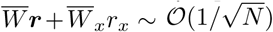 so that [28, 29, 43–45]

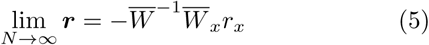

in the balanced state. Hence, the balanced state can only be realized when this solution has positi<u>ve</u> entries, *r*_*e*_, *r*_*i*_ *>* 0, which requires that [28, 29, 43] 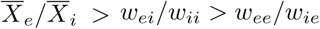.

Note that the derivation of Eq. (5) relied on an assumption that 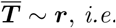, that the transfer of mean input to firing rates is 𝒪 (1). The analysis of correlations in balanced networks requires similar assumptions. Specifically, our analysis of correlations relies on an assumption that ⟨*T*_*a*_, *U*_*b*_⟩ ∼⟨*S*_*a*_, *U*_*b*_⟩ or

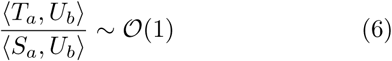

for *U* = *X, S* as *N* → ∞. A more precise and slightly weaker assumption that is sufficient for our analysis is that

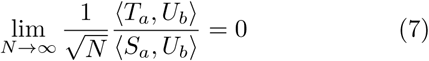

for *U* = *X, S*, which is implied by Eq. (6). Note that this does not imply that ⟨*S*_*a*_, *U*_*b*_ ⟩ cannot be much larger or smaller than ⟨*T*_*a*_, *U*_*b*_ ⟩, but only that their ratio does not converge to zero or diverge to ∞. as *N* → ∞. The validity of this assumption is discussed in more detail in Appendix A and the end of Section III.

Importantly, we do not need to know the value of the fraction in Eq. (6), the fraction does not need to converge to a limit as *N* → ∞., and ⟨*S*_*a*_, *U*_*b*_ ⟩ need not be linearly related to ⟨*T*_*a*_, *U*_*b*_, ⟩which contrasts to assumptions made by linear response theory.

## III. A REVIEW OF THE ASYNCHRONOUS BALANCED STATE

We first review previous work on correlated variability in balanced networks when spike trains in the external population are uncorrelated Poisson processes (Fig. 1A,B),

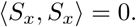

Since spike trains in the external population are uncorrelated, correlations between the external input to neurons in the recurrent network arise solely from overlapping feedforward synaptic projections with [31, 40, 46] (see Eq. (A1) in Appendix A for a derivation)

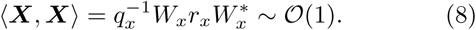

where 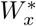 is the conjugate transpose of *W*_*x*_.

It would at first seem that this 𝒪 (1) external input correlation would lead to 𝒪 (1) correlations between neurons’ spike trains. In the asynchronous state, this is prevented by a cancellation between positive and negative sources of input correlation. In particular, correlations between neurons’ recurrent synaptic inputs, ⟨***R***, ***R***⟩, are also positive and 𝒪 (1), but these positive sources of input correlations are canceled by negative correlations between neurons’ recurrent and external inputs, ⟨***X***, ***R***⟩, in such a way that the total synaptic input correlation is weak,

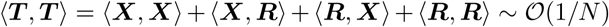

where ⟨***R***, ***X***⟩= ⟨***X***, ***R***⟩ *. In Appendix A, we review a derivation of spike train CSDs in the asynchronous state that gives

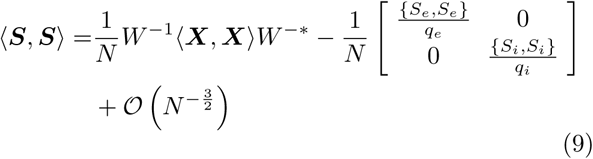

where {*S*_*b*_, *S*_*b*_}(*f)* is the average power spectral density of spike trains in population *b* = *e, i*.

Eq. (9) has been previously derived for the integrate-and-fire network models considered here (see equation S.22 in the supplement to [40]) and equivalent expressions have been derived for networks of binary neuron models (see equations 38-39 in the supplement to [31], equation 38 in [46], and equation 23 in [41]) as well as similar equations for similar models in other work [5, 14, 41, 47, 48]. In [46], the 𝒪 (*N* ^*-*3*/*2^) term is also derived for binary networks. These derivations have also been extended to spatially extended networks with distance-dependent connection probabilities and to networks with several sub-populations [40, 41]. The overall finding in this previous work is that spike train correlations in the recurrent network are 𝒪 (1*/N)*. Some exceptions to this 𝒪 (1*/N)* scaling have been demonstrated in spatially extended networks with distance-dependent connection probabilities and in networks with singular mean-field connectivity matrices [39–42, 46], a topic to which we return in Section VI.

To demonstrate these results, we first simulated a network of *N* = 10^4^ randomly and recurrently connected neurons receiving feedforward input from a population of *N*_*x*_ = 2000 uncorrelated Poisson-spiking neurons (Fig. 1A,B). As predicted, spiking activity in the recurrent network was asynchronous and irregular (Fig. 1C; mean spike count correlation between neurons with rates at least 1 Hz was 5.2×10^*-*4^) with approximate balance between external *(X)* and recurrent *(R)* synaptic input sources (Fig. 1D). Varying the network size, *N*, demonstrates the 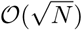 growth of mean external 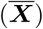 and recurrent 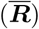 synaptic input current<u>s</u> that cancel to produce 𝒪 (1) mean total input current 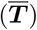 (Fig. 1E), as predicted by the mean-field theory of balance (Eq. (5) and surrounding discussion). As a result, firing rates converge toward the limiting values predicted by Eq. (5) (Fig. 1F).

As predicted by the analysis of the asynchronous state, the mean covariances between individual sources of synaptic inputs appear 𝒪 (1) (Fig. 1G), but cancel to produce total input covariance and spike count covariance that vanish as network size, *N*, increases (Fig. 1G–I).

Applying the approximation in Eq. (9) requires knowledge of neurons’ mean-field power spectral densities, {*S*_*e*_, *S*_*e*_} (*f)* and {*S*_*i*_, *S*_*i*_}(*f)*. While numerical methods for approximating power spectral densities in networks of integrate-and-fire neurons have been developed using Fokker-Planck techniques [8, 9, 11, 12, 49], these approaches require a diffusion approximation that is not justified for our model (see Discussion). Instead, we considered two ways to test Eq. (9) that do not require a diffusion approximation or Fokker-Planck techniques.

First, we noted that the first term, (1*/N)W* ^*-*1^ ⟨***X***, ***X***⟩*W* ^*-\**^, in Eq. (9) only involves known parameters and does not require knowledge of {*S*_*e*_, *S*_*e*_*}* or {*S*_*i*_, *S*_*i*_}. Therefore, we were able to directly compare the exact value of this term to spike count covariances computed from simulations, demonstrating a relatively close match (compare solid and dashed blue in Fig. 1I).

Next, we noted that the contribution of the second term in Eq. (9) can be estimated empirically from simulations by computing the average spike count variance in the recurrent network (since {*S*_*b*_, *S*_*b*_}(0) is proportional to spike count variance over large window sizes). This yields a semi-analytical approach to testing the accuracy of Eq. (9) with only the 𝒪 (*N* ^*-*3*/*2^) term ignored. This approach improved the approximation to mean inhibitory-inhibitory spike count covariances at large *N* (compare solid and dotted in Fig. 1I), but left some error even at *N* = 2 × 10^4^.

Previous work showed that intrinsically generated correlations can dominate in large balanced networks [46], but in our simulations external input correlations captured by the first term in Eq. (9) dominated over a range of *N* values. We conclude that the relative contribution of the three terms in Eq. (9) can depend on model details and parameters. This is clarified by noting that a sufficiently large reduction in the external input covariance, ⟨***X***, ***X*** ⟩, would necessarily prevent the first term in Eq. (9) from dominating because the first term is proportional to ⟨***X***, ***X***⟩, but the second term should remain non-zero even for vanishing ⟨***X***, ***X***⟩. At the extreme, when ⟨***X***, ***X*** ⟩= 0, the first term is zero, but the second term is non-zero for such networks due to intrinsically generated variability [46].

To demonstrate these ideas, we followed a procedure from [46] by varying *N*_*x*_ while scaling feedforward connection probabilities, *p*_*ex*_ and *p*_*ix*_, so as to keep the average number of external inputs, *p*_*ex*_*N*_*x*_ = *p*_*ix*_*N*_*x*_ = 200, fixed. All other parameters, including the size, *N*, of the recurrent network were kept fixed. In this scenario, an increase in *N*_*x*_ causes a decrease in ⟨***X***, ***X***⟩because it leads to fewer overlapping external inputs (as evidenced by the appearance of 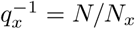 in Eq. (8); but note that *W*_*x*_ is fixed as we change *N*_*x*_). In addition, we simulated a version of the network in which each neuron received uncorrelated input from a private external population so that external input had the same univariate statistics as our previous simulations, but these external inputs were uncorrelated, ⟨***X***, ***X***⟩= 0. This represents the *N*_*x*_ *→* ∞ limit of the previous simulations. Finally, we simulated a deterministic version of the network in which external inputs were replaced by time-constant input, 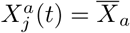, which also implies ⟨***X***, ***X***⟩= 0 with mean external input unchanged.

Examining these simulations, the magnitude of spike train covariance is non-monotonic with increasing *N*_*x*_ (Fig. 1J) because it changes sign and becomes increasingly negative at larger values of *N*_*x*_ (Fig. 1K). The first term in Eq. (9) dominates the other two terms (the dashed, solid, and dotted curves are close) for smaller values of *N*_*x*_ because external input covariance is larger in this regime. However, at larger values of *N*_*x*_, the first term no longer dominates (Fig. 1K, inset, the dashed curve is far from the solid curve) because external input covariance is small. Moreover, even accounting for the second term in Eq. (9) produces a somewhat inaccurate approximation when external input covariance is small (Fig. 1J,K, compare dotted to solid at larger *N*_*x*_), consistent with previous findings that the 𝒪 (*N* ^*-*3*/*2^) term can contribute significantly at large but finite *N* [46].

When external input is noisy but independent, simulations produce weak, but non-zero spike train correlations (Fig. 1J,K, filled circles) representing the *N*_*x*_ *→*∞ limit of the previous simulations [46]. In this regime, the first term in Eq. (9) is zero, but the second term produces a relatively accurate approximation to excitatory-excitatory and inhibitory-inhibitory spike count covariances (Fig. 1J,K, compare filled circles to asterisks), but note that the second term in Eq. (9) is zero for excitatory-inhibitory pairs. Similar results are observed when exte<u>rn</u>al input is deterministic and constant in time 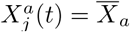; Fig. 1J,K, compare open circles to crosses), but spike count variances (and therefore the {*S*_*b*_, *S*_*b*_}(0) terms) are smaller when input is deterministic. Repeating the simulation from Fig. 1I,J with different connectivity parameters showed similar overall results (see Appendix B and Fig. 7A,B). See [5, 8–12, 14, 41, 43, 46, 47] for other studies of correlated variability in networks with uncorrelated external input.

So far we have focused on spike count covariance over large window sizes, which is proportional to low-frequency CSD. We next computed the full spike train CSDs from simulations with *N* = 10^4^ and *N*_*x*_ = 2000. The first two terms in Eq. (9) accurately capture most of the low-frequency CSD for these parameters (Fig. 1I–L), but simulations show a high-frequency peak in the mean CSD that is not captured by these terms (Fig. 1L).

To understand why Eq. (9) becomes inaccurate at higher frequencies for the network considered here, recall that the derivation of Eq. (9) relies on the assumptions made by Eqs. (6) and (7), which posit that neural transfer is 𝒪 (1). However, for EIF and many other neuron models, sufficiently high-frequency input fluctuations cannot be reliably be transferred due to the filtering imposed by membrane potential dynamics. More specifically,

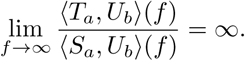

Hence, while Eqs. (6) and (7) might be true for any fixed *f*, the 𝒪 (1) term on the right side of Eq. (6) diverges with *f*. Analogously, when *f* is large, larger values of *N* must be considered before the ratio in Eq. (7) is close to zero. Mathematically speaking, the limit in Eq. (7) convergeces pointwise but not uniformly in *f*. This caveat is technically inconsequential for any fixed *f* in the *N →* ∞ limit analyzed in the derivation of Eq. (9), but all networks are finite. For any finite-sized network, the derivation of Eq. (9) is not justified at sufficiently high frequencies. In Appendix A, we show that for finite *N* our derivations are accurate at frequencies for which

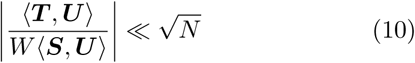

for ***U*** = ***X***, ***S*** where the division is applied element-wise to the matrices. As noted above, for any fixed *f*, this condition is satisfied in the *N →* ∞ limit, but for any fixed *N* the condition is violated at sufficiently large *f*.

The left-hand-side of Eq. (10) is difficult to compute for networks of integrate-and-fire neurons because we do not know the values of ⟨***T***, ***U***⟩ or ⟨***S***, ***U***⟩, but we next show that linear response theory can provide a useful rough approximation.

In the context of the model considered here, linear response theory is defined by making the approximation [6, 8, 50–55]

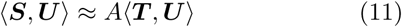

for *U* = *S, T, X* where

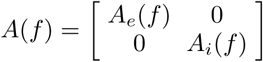

and *A*_*a*_(*f*) is the average “susceptibility function” of neurons in population *a* = *e, i*. Eq. (11) provides an accurate approximation of correlation transfer whenever correlations in the network are not too strong.

Combining Eq. (10) with Eq. (11) gives the condition

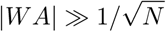

which can be written element-wise in terms of scalars as

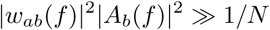

for *a, b* ∈ {*e, i*}. For the exponentially decaying synapses considered here, we have 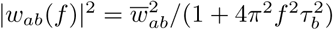 where 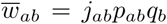 and *τ*_*b*_ is the synaptic time constant. However, we do not know the susceptibility functions, *A*_*b*_(*f*). Much like the power spectral densities, {*S*_*b*_, *S*_*b*_}(*f*), the susceptibility functions, *A*_*b*_(*f*), in networks of integrate-and-fire neurons are difficult to compute without a diffusion approximation that is not justified for our model (see Discussion). Due to the linearity of sub-threshold membrane dynamics, however, EIF neurons in a fluctuation dominated regime have susceptibility functions that are approximated by 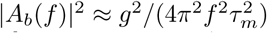 at moderately large *f* [52, 56, 57] where *τ*_*m*_ = *C*_*m*_*/g*_*L*_ is the membrane time constant and *g* = *A*_*b*_(0) is the gain of the neuron, *i.e*., the derivative of the neuron’s f-I curve. Putting this all together gives the condition

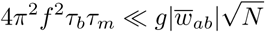

so we can expect that Eq. (9) is accurate for frequencies sufficiently lower than

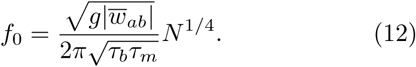

To obtain a conservative estimate that will be valid for all combinations of *a, b* ∈ {*e, i*}, we chose the smallest value of 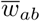 used in Fig. 2L, which is 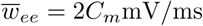 (where *C*_*m*_ is the arbitrary membrane capacitance), and the largest value of *τ*_*b*_, which is *τ*_*e*_ = 8ms. The membrane time constant for the neurons used in our simulations is *τ*_*m*_ = 15ms and *N* = 10^4^ in Fig. 1L. The only quantity missing from Eq. (12) is the gain, *g*. To obtain a rough estimate of *g*, we fit a rectified quadratic function to the relationship between<u>a</u>ll neurons’ firing rates and mean total inputs (*r*_*j*_ and 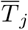), then computed the derivative of the fitted quadratic at the mean input value. The same approach was used in previous work [40, 58] to estimate a mean-field gain. In doing so, we obtained a gain approximation of *g* = 0.014(*C*_*m*_*mV*)^*-*1^. Combining these values gives an approximate cutoff frequency of *f*_0_ = 24Hz. Indeed, Fig. 1L shows that Eq. (9) starts to become inaccurate somewhere just below this value of *f*, particularly for excitatory-excitatory (red) and excitatory-inhibitory (purple) neuron pairs.

**FIG. 2.**
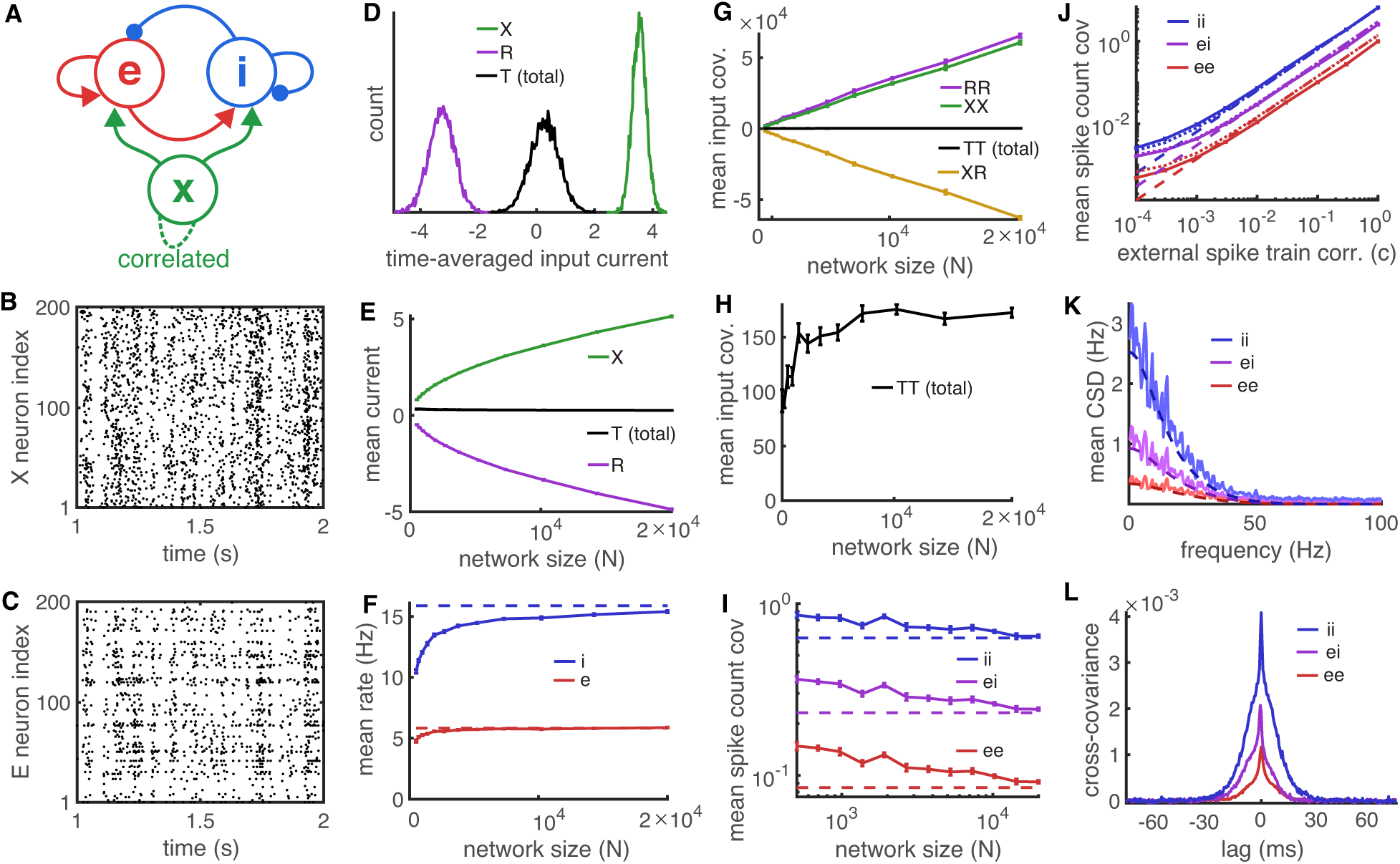
The correlated state in densely connected balanced networks. A–I and K) Same as Fig. 1A–I and L respectively except spike trains in the external population, *X*, were correlated Poisson processes with spike count correlation *c* = 0.1 and Eq. (19) was used for the dashed lines in panels I and K. **J)** Mean spike count covariance between neurons in the recurrent network as a function of the correlation, *c*, between spike trains in the external population. Dashed curves are from Eq. (19) and dotted are from Eq. (9). **L)** Mean cross-covariance functions between neurons in the recurrent network (units ms^*-*2^; *N* = 10^4^). Synaptic time constants were *τ* _*e*_ = 8, *τ* _*i*_ = 4, and *τ* _*x*_ = 10ms in all simulations and the correlation time constant for the spike trains in the external population was *τ* _*c*_ = 5ms. In all simulations, recurrent and feedforward connection probabilities are all *p*_*ab*_ = 0.1 for *a* = *e, i* and *b* = *e, i, x*.

In summary, when spike trains in the external population are uncorrelated, the first two terms in Eq. (9) give an accurate approximation to spike count covariances over long time windows (equivalently, low-frequency CSDs), but the second term can be difficult to compute directly and the equation loses accuracy when evaluating the CSD at higher frequencies. We next extend these results to networks in which spike trains in the external population are correlated.

## IV. THE CORRELATED STATE IN BALANCED NETWORKS

Above, we reviewed the asynchronous state in which uncorrelated spike trains in the external layer, ⟨*S*_*x*_, *S*_*x*_⟩= 0, produce moderate external input covariance, ⟨***X***, ***X***⟩ ∼ 𝒪 (1), and weak spike train correlations, ⟨***S***, ***S***⟩ ∼ 𝒪 (1*/N*). We next show that moderate correlations between spike trains in the external population (Fig. 2A),

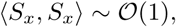

leads to large covariance between neurons’ external inputs, ⟨***X***, ***X***⟩ ∼ 𝒪 (*N*), and moderate correlations between spike trains in the recurrent network, ***S***, ***S*** ∼ 𝒪 (1).

We outline the derivation of correlations in such networks here and give a more detailed derivation in Appendix A. In addition to the assumption made by Eqs. (6) and (7), all of our derivations follow from a few simple arithmetical rules that rely on the bilinearity of the operator ⟨*·, ·*⟩. Specifically,

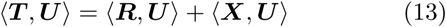

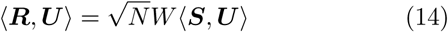

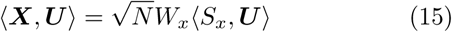

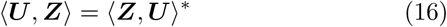

for any ***U, Z*** = ***E, I, X, R, S***, *S*_*x*_, ***T*** where ***A***^*\**^ is the conjugate-transpose of ***A*** and where we omit smaller order terms here and below (the derivations in Appendix A keep track of these terms). Eq. (13) follows from the fact that total input is composed of recurrent and external sources, ***T*** = ***R*** + ***X***. Eqs. (14) and (15) follow from the fact that recurrent and external inputs are composed of linear combinations of 𝒪 (*N*) spike trains, *c.f*. Eq. (1), and that synaptic weights are 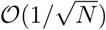. Eq. (16) is simply a property of the Hermitian cross-spectral operator.

We first derive the CSD between external inputs to neurons in the recurrent network. Applying Eqs. (15) and (16) gives

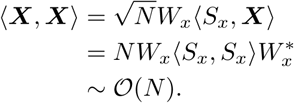

Hence, 𝒪 (1) covariance between the spike trains in the external population induces 𝒪 (*N*) covariance between the external input currents to neurons in the recurrent network. This is a result of the effects of “pooling” on covariances, namely that the covariance between two sums of *N* correlated random variables is typically 𝒪 (*N*) times larger than the covariances between the individual summed variables [31, 59, 60].

We next derive the CSD between spike trains and external inputs. First note that

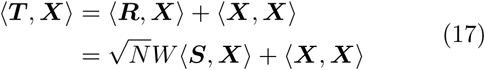

from Eqs. (13) and (14). It follows from our assumption that neuronal transfer is 𝒪 (1) (see Eq. (6)) that (***T, X***) ∼⟨***S***, ***X***⟩ which, combined with Eq. (17), gives

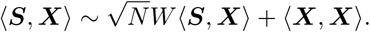

Because ⟨*S*, ***X***⟩ appears on both sides of this equation with a 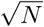 coefficient on one side and not the other, this is only consistent if there is a cancellation between the two terms on the right hand side. Specifically, this cancellation implies that

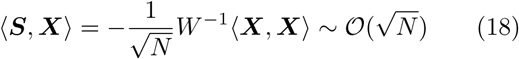

since ⟨***X***, ***X***⟩ ∼*O*(*N)*. We can now calculate the CSD between spike trains in the recurrent network. First note that

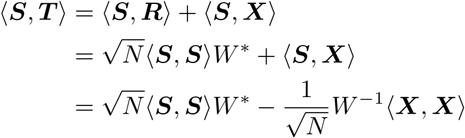

which follows from Eqs. (13), (14), and (18). The assumption of 𝒪 (1) transfer from Eq. (6) implies that

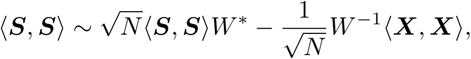

which is only consistent if there is cancellation between the terms on the right hand side. This cancellation can only be realized if

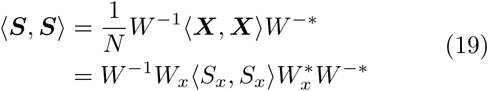

which is 𝒪 (1) and where we have omitted terms of smaller order.

In summary, 𝒪 (1) covariance between spike trains in the external population produces 𝒪 (*N)* covariance between neurons’ external inputs, but 𝒪 (1) covariance between spike trains in the recurrent network on average. We hereafter refer to this state as the “correlated state” since it produces moderately strong spike train correlations in contrast to the asynchronous state characterized by extremely weak spike train correlations. The reduction from 𝒪 (*N)* external input covariance to *𝒪* (1) spike train covariance arises from the same cancellation mechanism that reduces 𝒪 (1) external input correlation to *𝒪* (1*/N)* spike train correlations in the asynchronous state.

Notably, evaluating Eq. (19) does not require knowledge of neurons’ power spectral densities or spike count variances, but only depends on synaptic parameters and external spike train statistics. This is beneficial because the direct numerical computation of power spectral densities or spike count variances in networks of integrate-and-fire neurons can be difficult (see discussion in Section III). This contrasts to Eq. (9) that depends on mean power spectral densities or spike count variance through the second term. Note that Eq. (9) is also valid in the correlated state (see Appendix A). Specifically, Eq. (19) can be obtained from Eq. (9) by taking ***⟨X, X⟩*∼** (*N)* and ignoring smaller order terms. Hence, Eq. (9) is more accurate at finite *N*, but requires knowledge of mean power spectral densities whereas Eq. (19) is accurate as *N →* ∞ in the correlated state.

To demonstrate these results, we simulated a network of *N* = 10^4^ neurons identical to the network from Fig. 1 except that spike trains in the external population were correlated Poisson processes (Fig. 2A,B) with

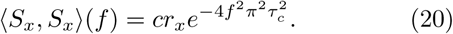

Here, *r*_*x*_ = 10Hz is the same firing rate used in Fig. 1, *c* = 0.1 quantifies the spike count correlation coefficient between the spike trains in the external population over large counting windows, and *τ*_*c*_ =5ms quantifies the timescale over which these correlations occur. See Appendix C for a description of the algorithm used to generate the spike trains.

The recurrent network exhibited moderately correlated spike trains in contrast to spike trains in the asynchronous state (Fig. 2C, compare to Fig. 1C; mean spike count correlation between neurons with rates at least 1 Hz was 0.077 in the correlated state). As in the asynchronous state, external and recurrent synaptic input sources approximately canceled (Fig. 2D), as predicted by balanced network theory. Varying *N* demonstrates that <u>t</u>he network exhibits the same cancellation between 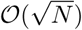 mean external and recurrent synaptic input sources and that Eq. (5) for the mean firing rates is accurate (Fig. 2E,F).

As predicted by the analysis of the correlated state, the covariance between individual sources of input currents appear 𝒪 (*N)* (Fig. 2G), but cancel to produce much smaller, approximately 𝒪 (1), total input covariance (Fig. 2G,H). Mean spike count covariances also appear 𝒪 (1) and converge toward the limit predicted by Eq. (19) (Fig. 2I). The timescale of correlations between neurons in the recurrent network is quantified by their mean cross-covariance functions (Fig. 2L). Repeating the simulation from Fig. 2I with different connectivity parameters showed similar overall results (see Appendix B and Fig. 7C).

We next investigated the dependence of spike count covariance in the recurrent network on the magnitude, *c*, of spike train correlations in the external population. The correlated state is characterized by *c* ≠ 0 and the asynchronous state by *c* = 0. This transition is continuous in the sense that, for fixed *N*, sufficiently small values of *c >* 0 generate correlations similar to those in the asynchronous state. However, the transition between asynchronous and correlated spiking with increasing *c* becomes more abrupt for larger values of *N*. Specifically (see Appendix A),

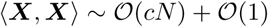

and

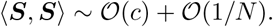

Hence, when *N* is large, even very small values of *c ≫* 1*/N* should produce much stronger correlations than those produced in the asynchronous state (*c* = 0).

Indeed, simulations in which *N* = 10^4^ is fixed and *c* is varied between *c* = 10^*-*4^ and *c* = 1 show that Eq. (19) is relatively accurate for *c* larger than 10^*-*3^ (Fig. 2J, compare solid to dashed). When *c* is so small that Eq. (19) is inaccurate, Eq. (9) provides a more accurate approximation (Fig. 2J, compare solid to dotted), but requires the empirical estimation of power spectral densities. The approximate linear relationship between spike train correlation and external input correlation (at larger values of *c ≫*1*/N)* is consistent with data from cultured cortical populations [30], but those data also show moderate spike train correlations in the absence of external input correlation, in contrast to our model. These could arise from pattern forming intrinsic dynamics (see Discussion).

Mean CSDs between spike trains in the recurrent network closely matched the theoretical predictions from Eq. (19) over a range of frequencies (Fig. 2K) even though our theory only strictly applies to CSDs at sufficiently low frequencies (see Eq. (10) and surrounding discussion). To understand why this is the case, we again turn to linear response theory. Under the assumption that neurons’ input CSDs are approximately linearly transferred to their spike train CSDs (*i.e*., Eq. (11)) one can derive an approximation to spike train CSDs (see [8–14, 41, 46] for derivations of this and similar approximations)

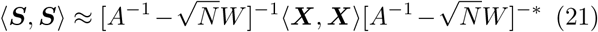

where recall that *A*(*f)* is the matrix of mean-field susceptibility functions. In the *N* limit for any fixed *f*, Eq. (21) reduces to Eq. (19). At finite *N* → ∞, the tw<u>o</u> equations are approximately equivalent whenever 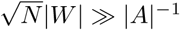. Note that this is equivalent to the condition in Eq. (10) and the surrounding discussion, where it was argued that this condition is violated for sufficiently large *f* because *|A*^*-*1^(*f)|* → ∞ as *f* → ∞. Therefore, when *f* is suf<u>fic</u>iently large, 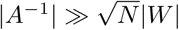 and we can ignore the 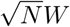 terms in Eq. (21) to get

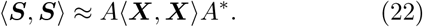

for sufficiently large *f*. In other words, at frequencies too high (timescales too fast) for synapses to track, external input CSD is transferred directly to spike train CSD through the neurons’ susceptibility functions and recurrent connectivity plays a vanishing role. For the simulations in Fig. 2K, 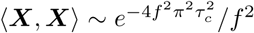 at large *N* and *f*, but as discussed in Section III, |*A*| ^2^ ∼1*/f* ^2^ for EIF neurons in the fluctuation dominated regime. Therefore, ⟨***X***, ***X*** ⟩ decays to zero much faster than *A* for large *f*. Hence, for the larger values of *f* for which Eq. (22) is accurate, *A* ⟨***X***, ***X*** ⟩ *A*\*≈ 0. This explains why the approximation in Eq. (19) does not lose accuracy at higher frequencies in Fig. 2K: because ⟨***S***, ***S***⟩ ≈ 0 anyways for larger values of *f*.

## V. THE CORRELATED STATE PRODUCES TIGHT BALANCE BETWEEN EXCITATORY AND INHIBITORY INPUT FLUCTUATIONS CONSISTENT WITH CORTICAL RECORDINGS

We have so far considered cancellation between positive and negative sources of input correlations at the mean-field level, *i.e*., averaged over pairs of postsynaptic neurons (Figs. 1G,H and 2G,H). Previously published *in vivo* intracellular recordings from neurons in rat barrel cortex in reveal that this cancellation occurs even at the level of single postsynaptic neuron pairs [23]. Specifically, paired intracellular recordings of spontaneous neural activity were performed between nearby neurons (distance < 500 *µ*m) in the barrel cortex of lightly anesthetized rats in current-clamp mode. When one neuron was clamped near its inhibitory reversal potential and another neuron is clamped near its excitatory reversal potential (spiking suppressed with QX-314), recorded membrane potential fluctuations are approximately mirror images of one another (Fig. 3A, top). Similarly, if both neurons are held near their excitatory reversal potential (Fig. 3A, middle) or both near their inhibitory reversal potential (Fig. 3A, bottom), recorded membrane potential fluctuations are highly correlated. This implies that fluctuations in the excitatory and inhibitory synaptic input to one neuron are strongly correlated with fluctuations in the excitatory and inhibitory input to other nearby neurons (see [23] for more details and interpretation).

**FIG. 3.**
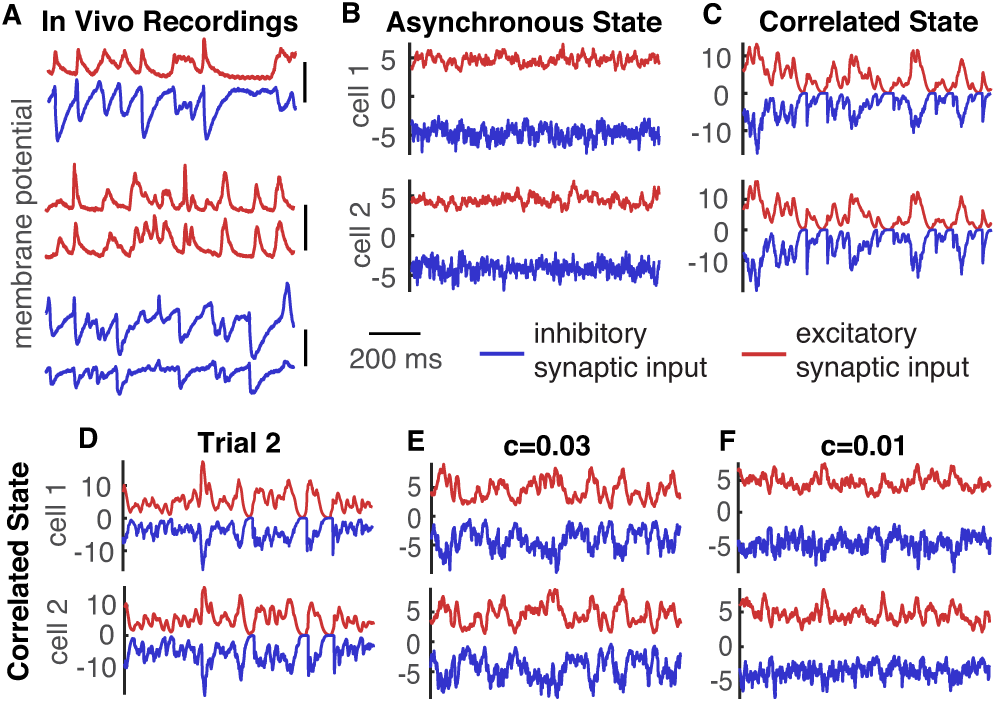
Excitatory-inhibitory tracking in vivo and in simulations. **A)** In vivo membrane potential recordings from neurons in rat barrel cortex, reproduced from [23]. Each pair of traces are simultaneously recorded membrane potentials. Red traces were recorded in current clamp mode near the reversal potential of inhibition and blue traces near the reversal potential of excitation (with action potentials pharmacologically suppressed), so red traces are approximately proportional to excitatory input current fluctuations and blue traces approximate inhibitory input current fluctuations. Vertical scale bars are 20mV. For ease of comparison with the current traces in panels B–F, the red voltage trace in the top of A was plotted above the corresponding blue trace. **B,C)** Excitatory (red) and inhibitory (blue) synaptic input currents to two randomly selected excitatory neurons in the asynchronous (B) and correlated (C) states. Simulations were the same as those in Figs. 1B-D and 2A-D respectively. **D)** Same as C, but for a second trial with the same connection matrix. **E,F)** Same as C, but correlations between the external population’s spike trains changed to *c* = 0.03 and 0.01 respectively (from *c* = 0 in B and *c* = 0.1 in C).

To test whether this phenomenon occurred in our simulations, we randomly chose two neurons and decomposed their synaptic input into the total excitatory (*E* + *X*) and the inhibitory (*I*) components. In the asynchronous state, input current fluctuations were fast and largely unshared between neurons or between current sources in the same neuron (Fig. 3B), in contrast to evidence from *in vivo* recordings. Input current fluctuations in the correlated state were larger and highly synchronized between neurons (Fig. 3C), consistent with evidence from *in vivo* recordings. This precise tracking of fluctuations in excitatory and inhibitory synaptic currents is referred to as “tight balance” [61] (as opposed to the “loose balance” of the asynchronous state). The results would be similar if we decomposed inputs into their external (*X*) and recurrent (*R* = *E* + *I*) sources instead of excitatory (*E* + *X*) and inhibitory (*I*). The large fluctuations in synaptic currents in the correlated state are shared between neurons, but exhibit trial-to-trial variability (compare Fig. 3C to D), clarifying that they represent noise correlations instead of signal correlations.

An intuitive way of understanding the tight balance produced in the correlated state is to note that shared fluctuations in neurons’ external synaptic input is 𝒪 (*cN)*+ 𝒪 (1) and these fluctuations are inherited by the recurrent network so that fluctuations between neurons’ excitatory (*E* + *X*) or inhibitory (*I*) synaptic inputs are also 𝒪 (*cN)* + 𝒪 (1). Hence, shared fluctuations are 𝒪 (*N)* in the correlated state (*c* > 0) and 𝒪 (1) in the asynchronous state (*c* = 0). Since connectivity in the networks is homogeneous, the strongly correlated input current fluctuations also have a similar magnitude in the correlated state, so they appear to track each other. In the asynchronous state, the more weakly correlated fluctuations are “washed out” by uncorrelated variability in the network, so currents do not appear to closely track each other. This implies that the transition from the loose balance of the asynchronous state to the tight balance of the correlated state occurs continuously as *c* is increased from zero, but also that tight balance is realized at small values of *c* > 0 whenever *N* is large (compare to the discussion of Fig. 2J above). Indeed, simulations demonstrate this continuous transition between tight and loose balance as *c* is modulated (Fig. 3B,C,E,F).

Despite the striking differences between excitatory and inhibitory synaptic currents in the correlated versus asynchronous states, the distributions of spike count covariances were qualitatively somewhat similar in the two states (Fig. 4A). It is more common in the literature to report spike count correlations instead of spike count covariances and it is common to omit neurons with low firing rates. Therefore, we next computed the distribution of spike count correlations between excitatory neurons with firing rates of at least 1 Hz (Fig. 4B). It is easier to distinguish between the correlated and asynchronous states from these distributions, but the distributions are still somewhat similar. Note especially that, despite the differences between the excitatory and inhibitory synaptic currents between the *c* = 0 and *c* = 0.03 cases (Fig. 3B,E), the distributions of spike count correlations are qualitatively similar (Fig. 4A,B, compare light and dark gray). The mean spike count correlations differed by orders of magnitude across the two states (mean correlation between excitatory neurons’ spike trains 2.6 × 10^*-*4^, 2.4 × 10^*-*2^, 6.6 × 10^*-*2^ for *c* = 0, 0.03, 0.1 respectively) while the standard deviation of correlations was similar across the states (7.4 × 10^*-*2^, 8.1 × 10^*-*2^, and 1.2 × 10^*-*1^ for *c* = 0, 0.03, 0.1 respectively). Hence, while the distributions of correlations are qualitatively somewhat similar, the asynchronous state is distinguished by having a much larger standard deviation than mean correlation value [31, 48].

**FIG. 4.**
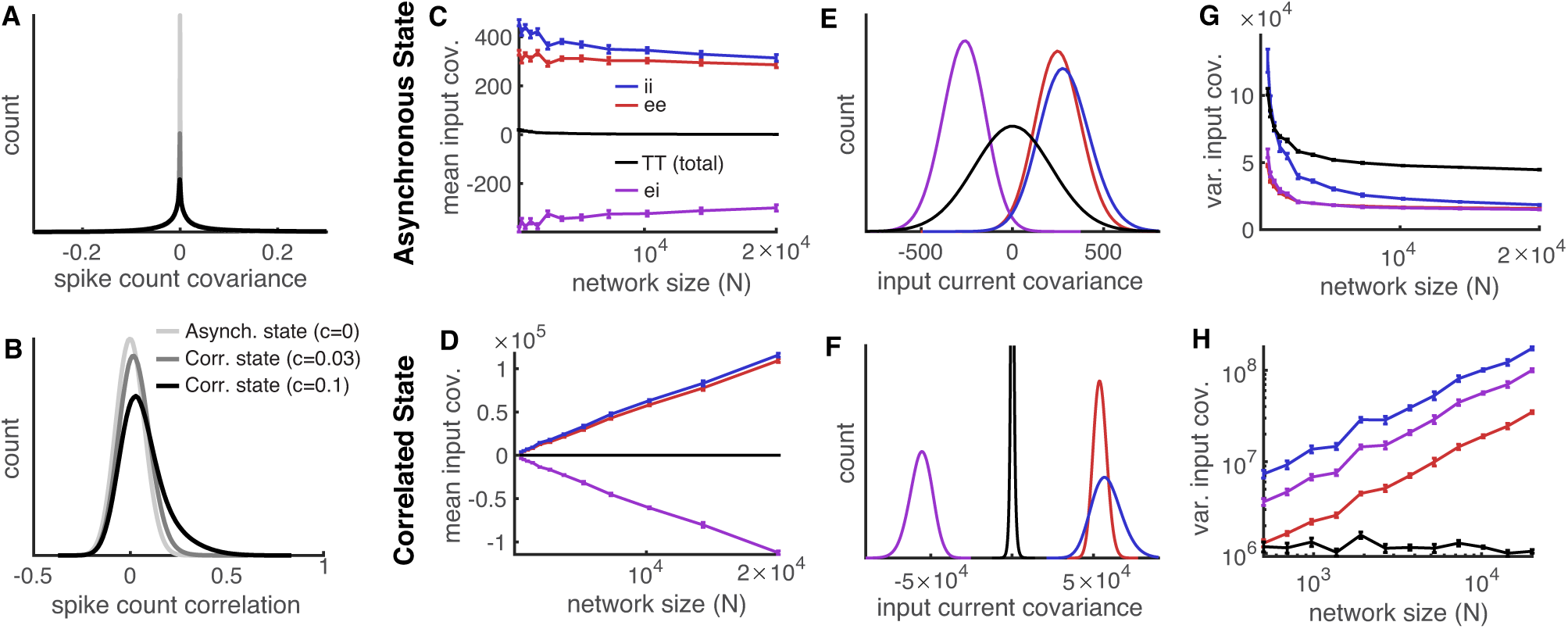
The scaling of mean and variance of excitatory and inhibitory input covariance in the asynchronous and correlated states. **A,B)** Distributions of spike count covariances (A) and correlations (B) between excitatory neurons in the asynchronous state (*c* = 0, light gray) and in the correlated state (*c* = 0.03, dark gray; *c* = 0.1, black). For spike count correlations, neurons with firing rates less than 1 Hz were omitted from the analysis. **C,D)** Same as Figs. 1G and 2G, except inputs were decomposed into their excitatory (*E* + *X*), and inhibitory (*I*) components instead of external and recurrent. Red curves show mean excitatory-excitatory input covariance, blue show inhibitory-inhibitory, purple show excitatory-inhibitory, and black curves show total (same as black curves in Figs. 1G and 2G). **E,F)** Histogram of input current covariances across all excitatory cell pairs for a network of size *N* = 10^4^. **G,H)** Same as C,D except we plotted the variance of covariances across cell pairs instead of the mean. As above, integrated currents have units *C*_*m*_*mV*, so input covariances have units 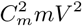 and the variance of covariances have units 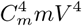 where *C*_*m*_ is the arbitrary membrane capacitance. In panels D,F,H we set *c* = 0.1.

To further quantify differences between covariances in the asynchronous and correlated states, we first computed the average covariance between the excitatory and inhibitory input to pairs of (excitatory) neurons in the network. These averages have the same dependence on network size, *N*, as they do when input currents are broken into external and recurrent sources (compare Fig. 4C,D to Figs. 1G and 2G). Specifically, in the asynchronous state, covariances between individual current sources are 𝒪 (1) on average, but cancel to produce weak 𝒪 (1*/N)* covariance between the total synaptic input to neurons on average (Fig. 4C). In the correlated state, the average covariance between individual input sources is 𝒪 (*N)* and cancellation produces 𝒪 (1) average total input covariance (Fig. 4D).

Hence, despite the precise cancellation of positive and negative sources of input covariance at the mean-field level in the asynchronous state (Fig. 4C), the tracking suggested by this cancellation is apparently not observed at the level of individual neuron pairs (Fig. 3C). To see why this is the case, we computed the distribution of input current covariances across all pairs of excitatory neurons. We found that these distributions were broad and the distribution of total input covariance was highly overlapping with the distributions of individual input current sources (Fig. 4E, the black distribution overlaps with the others). This implies that cancellation does not reliably occur at the level of individual pairs since, for example, the total input covariance for a pair of neurons can be similar in magnitude or even larger than the covariance between those neurons’ excitatory inputs.

The distributions of input covariances were strikingly different in the correlated state. The distribution of total input covariances was far narrower than the distributions of individual input current sources and the distributions were virtually non-overlapping (Fig. 4F). Hence, a precise cancellation between positive and negative sources of input covariance must occur for every neuron pair, leading to the tight balance observed in Fig. 3E.

These results are better understood by computing the empirical variance of input covariances across neuron pairs in simulations as *N* is varied. In the asynchronous state, the empirical variance of input covariances from all sources appear to scale like 𝒪 (1) (Fig. 4G). Since the mean input covariance between individual sources are also 𝒪 (1) (Fig. 4C), the overlap between distributions in Fig. 4E is expected. In the correlated state, the empirical variances of input covariances appear to scale like 𝒪 (*N)* except for the variance of the total input covariance, which appears to scale like 𝒪 (1) (Fig. 4H). If the variances scale like 𝒪 (*N)*, then the standard deviations would scale like 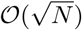. This, combined with the fact that the mean input covariances between individual sources scale like (*N)*, implies that the distributions in Fig. 4G will be non-overlapping when *N* is large. The same conclusions would be reached if we decomposed inputs into their external (*X*) and recurrent (*R* = *E* + *I*) sources instead of total excitatory (*X* + *E*) and inhibitory (*I*). Note, that the scaling of the variance of covariances reported here was only computed empirically and we have not derived these scalings analytically. It is possible that, at larger *N*, the scaling becomes different from how it appears from our simulations.

## VI. CORRELATED VARIABILITY FROM SINGULAR MEAN-FIELD CONNECTIVITY STRUCTURE

We have shown that 𝒪 (1) spike train correlations can be obtained in balanced networks by including correlations between neurons in an external layer (⟨*S*_*x*_, *S*_*x*_⟩ 𝒪 (1)), defining what we refer to as the “correlated state.” Previous work shows that 𝒪 (1) spike train correlations can be obtained in the recurrent network with uncorrelated external spike trains (⟨*S*_*x*_, *S*_*x*_ ⟩= 0) when the mean-field connectivity matrix is constructed in such a way that the recurrent network cannot achieve the cancellation required for these states to be realized [39–41]. This can be achieved using networks with several discrete subpopulations or networks with distance-dependent connectivity. We first review these previous results by considering networks with discrete sub-populations, then show that excitatory-inhibitory tracking is similar to the asynchronous state in this case.

The recurrent networks considered above have two statistically homogeneous sub-populations: one excitatory and one inhibitory and the external population is a single homogeneous population. Suppose instead that there are *K* sub-populations in the recurrent network, with the *k*th population containing *N*_*k*_ = *q*_*k*_*N* neurons where Σ*k q*_*k*_ = 1. Connectivity is random with *p*_*jk*_ denoting the co<u>nn</u>ection probability from population *k* to *j*, and 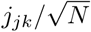 denoting the strengths of the connections for *j, k* = 1, *… K*. All neurons in population *k* are assumed to have the PSC kernel *η*_*k*_(*t*) which is again assumed to have integral 1. Similarly, suppose that the external network contains *K*_*x*_ sub-populations each with 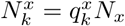 neurons where 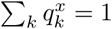. Feedforward connection probabilities and strengths are given by 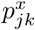 and 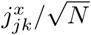 for *j* = 1, *… K* and *k* = 1, *…, K*_*x*_. Assume that 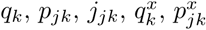 and 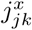 are all 𝒪 (1). We then define the *K* × *K* mean-field recurrent connectivity matrix by 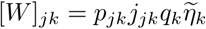 and the mean-field feedforward connectivity matrix by 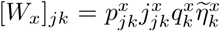. For all of the networks considered above, we had *K* = 2 and *K*_*x*_ = 1.

When *W* is an invertible matrix, Eqs. (5), (9), and (19) are equally valid for networks with several subpopulations as they are for the simpler networks considered above. Hence, the mean-field theory of firing rates and correlations extends naturally to networks with several populations [40, 41, 43–45, 58]. However, when *W* is singular, Eqs. (5), (9), and (19) cannot be evaluated. Instead, Eq. (5) can be re-written as

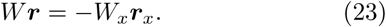

When *W* is singular, this equation only has a solution, ***r***, when 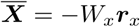 is in the range or “column space” of *W*. Otherwise, balance is broken. An in-depth analysis of firing rates in such networks is provided in previous work [44, 45, 58] (and extended to spatially continuo<u>us</u> networks in [41, 43, 58]), so we hereafter assume that 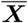 is in the range of *W* and balance is achieved.

A similar analysis may be applied to spike train CSDs. For simplicity, we assume here that spike trains in the external population are uncorrelated, ⟨ *S*_*x*_, *S*_*x*_ ⟩ = 0, since this is the case considered in previous work and since this is the case in which a singular *W* breaks the asynchronous state. Eq. (9) can be re-written as

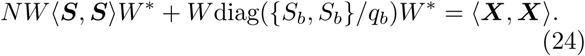

where diag({*S*_*b*_, *S*_*b*_}/*q*_*b*_) is a diagonal matrix with ratio of the mean power spectral density, {*S*_*b*_, *S*_*b*_}, divided by *q*_*b*_ = *N*_*b*_/*N* on the diagonal, and where we have ignored smaller order terms. When *W* is singular, Eq. (24) is not guaranteed to have a solution, ⟨**⟨ *S, S* ⟩ ⟩**. More precisely, a solution can only exist when the *K* × *K* matrix, ⟨ ***X, X* ⟩** ⟩, is in the range of the linear operator ℒ defined by

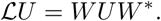

In that case, Eq. (24) has a solution so that ⟨**⟨ *S, S* ⟩**⟩ ∼ *𝒪𝒪* (1/*N)* and the asynchronous state is still realized. However, if ⟨ ***X, X* ⟩** ⟩ is not in the range of ℒ, the asynchronous state cannot be realized because Eq. (24) does not have a solution.

Using Eqs. (13), (14), and (16), we can write the meanfield total input CSD as

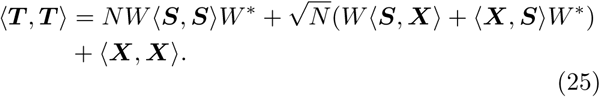

If *W* is not invertible, then *W\** has a non-trivial nullspace. Let *v*_1_, *v*_2_, *…, v*_*n*_ be a basis for the nullspace of *W\** and define

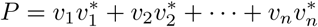

which is a self-adjoint matrix that defines the orthogonal projection onto the nullspace of *W \**. Note that *P* is a Hermitian matrix (*P* = *P \**) and *PW* = *W *P* = 0 (the zero matrix). Define the projection *A*_0_ = *P AP* for any matrix *A*. Unless ⟨ ***X, X* ⟩** is carefully constructed otherwise, we can expect that

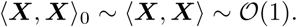

Then take the projection of both sides of Eq. (25) above to get

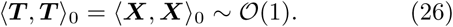

Hence, the total input CSD is 𝒪𝒪 𝒪(1) when **⟨*X, X* ⟩** is not in the range of ℒ, even though it is **⟨*X, X* ⟩** /*N* when *W* is invertible (*i.e*., in the asynchronous state). Moreover, the structure of **⟨ *T, T* ⟩** is given to highest order in *N* by **⟨*X, X* ⟩** _0_ = *P* **⟨*X, X* ⟩** *P*, which can be computed exactly from knowledge of the structure of **⟨*X, X* ⟩** and *W*.

When neural transfer from ***T*** to ***S*** is 𝒪𝒪 𝒪(1) (see Eq. (6) and surrounding discussion), this implies that **⟨ *S, S* ⟩ ∼** 𝒪𝒪 𝒪(1)so that the asynchronous state is broken when **⟨*X, X* ⟩** is not in the range of ℒ. While we cannot be certain that **⟨ *S, S* ⟩** has the same structure as **⟨ *T, T* ⟩,** it should have a similar structure as long as neural transfer of correlations is similar for each sub-population.

Previous work [41] derived similar conditions on the cancellation required for realizing the asynchronous state in networks of binary neurons, extended the analysis to spatially extended networks, and derived analytical expressions for covariances in these networks. Other work [39, 40] analyzed singular connectivity in networks of integrate-and-fire neurons and the extension to spatially extended networks, but only showed when the asynchronous state was broken and did not derive the structure of covariances when it was broken.

To demonstrate these results, we consider the same network from above with re-wired feedforward projections from the external population. Specifically, divide the excitatory, inhibitory, and external populations each into two equal-sized sub-populations, labeled *e*_1_, *i*_1_, *x*_1_, *e*_2_, *i*_2_, and *x*_2_ where population *a*_*k*_ contains *N*_*a*_/2 neurons. Hence the network has the same total number of neurons as before, but we have simply sub-divided the populations. To distinguish this network from the one considered in Figs. 1 and 2, we refer to the previous network as the 3-population network and to this modified network as the 6-population network.

We re-wire the feedforward connections so that *x*_1_ only connects to *e*_1_ and *i*_1_, and *x*_2_ only projects to *e*_2_ and *i*_2_. Specifically, we set the connection probabilities to 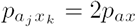 if *j* = *k* and 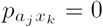 if *j ≠ k* for *a, b* = *e, i* and *j, k* = 1, 2, where *p*_*ab*_ are the connection probabilities for the 3-population network and 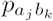 for the 6population network. This re-wiring assures that neurons in the recurrent network receive the same number of feedforward connections on average from the external population. The recurrent connectivity structure is not changed at all. Specifically, we set 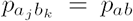 for *a, b* = *e, i*. All connection strengths are unchanged, 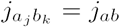 for *a* = *e, i* and *b* = *e, i, x* and all neurons in the external population have the same firing rate, *r*_*x*_, as before. See Fig. 5A for a schematic of this network.

**FIG. 5.**
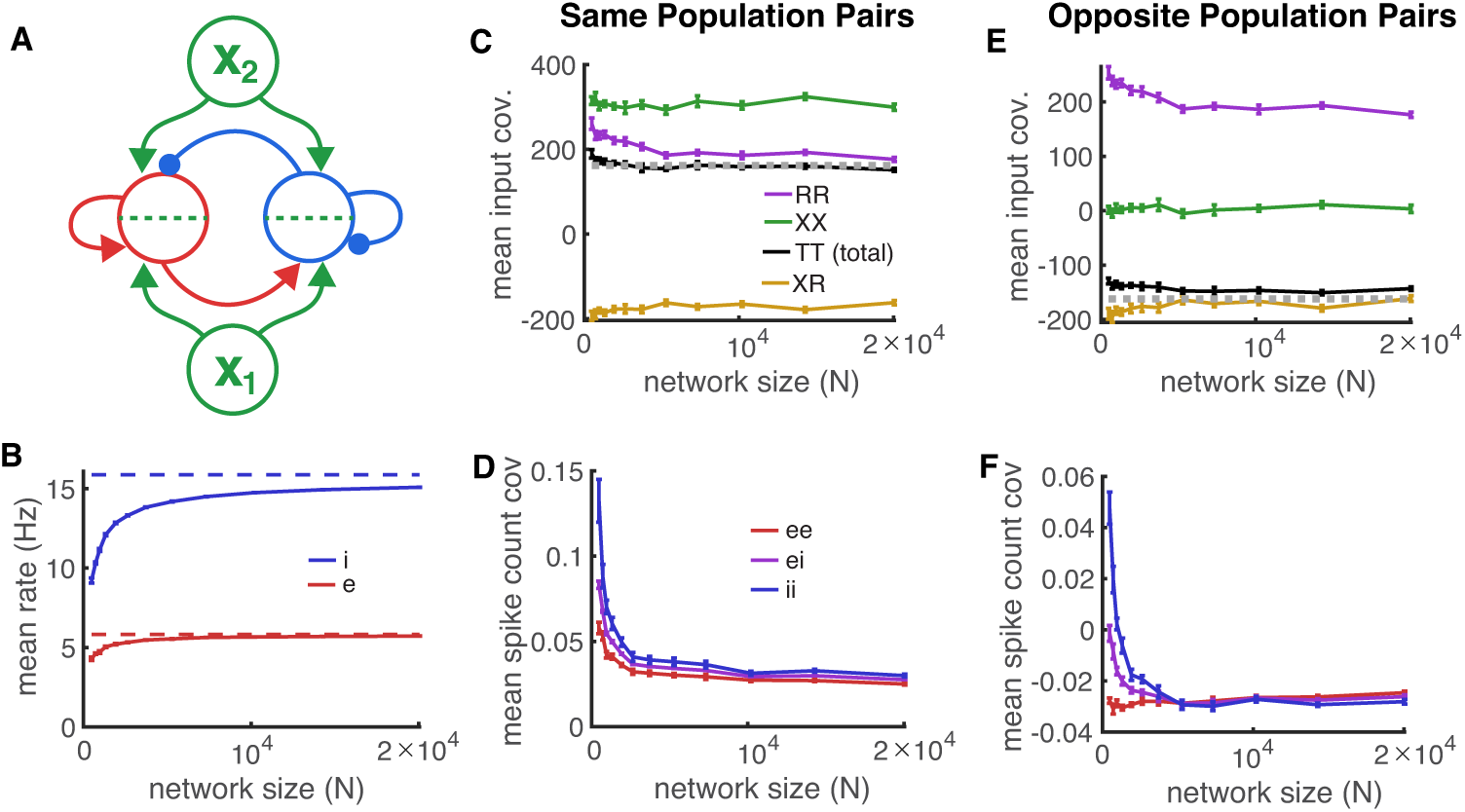
Correlated variability in a balanced network with singular mean-field connectivity matrix. **A)** Network schematic. The recurrent network is statistically identical to the networks considered previously, but there are two external populations that each connect to a different half of the neurons in the recurrent network. **B)** Same as Fig. 1F, but for the multi-population network from A. **C)** Same as Fig. 1G, but for the network in A and where input covariances are only averaged over postsynaptic neurons in the same group (both postsynaptic cells in *e*_1_ or both in *e*_2_). The dashed gray curve shows the theoretical prediction for total input covariance (the black curve) from Eq. (27). **D)** Same as Fig. 1I, but for the network in A and where spike count covariances are only averaged over postsynaptic neurons in the same group (first cell in *aj* and second cell in *bj* for *a, b* = *e, i* and *j* = 1, 2). **E,F)** Same as C and D, but covariances are computed between cells in opposite groups (one cell in *a*_1_ and the other cell in *b*_2_).

The feedforward mean-field connectivity matrix can be written in block form as

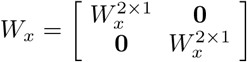

where **0** is the 2 × 1 zero-matrix and 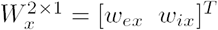 is the 2 × 1 feedforward connectivity matrix for the 3 population network. Note that *W*_*x*_ is 4 × 2 since there are 4 populations in the recurrent network and 2 populations in the external population. The recurrent mean-field connectivity matrix is

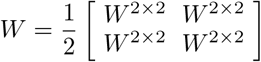

where

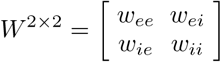

is the 2 ×2 recurrent connectivity matrix for the 3population network. Note that *W* is 4× 4. Here, *w*_*ab*_ = *p*_*ab*_*j*_*ab*_*q*_*b*_*ῆ*_*b*_(*f)* are the same values used above for analyzing the 3-population network.

Even though *W* is non-invertible, 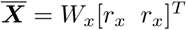 is in the range of *W* for this example, so firing rates in the balanced state can be computed using Eq. (23), and are identical to the firing rates for the 3-population networks considered above.

The nullspace of *W \** is spanned by the orthonormal vectors

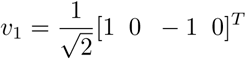

and

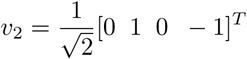

so the projection matrix is given in block form by

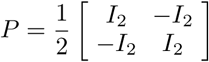

where *I*_2_ is the 2 ×2 identity matrix.

The external input CSD is determined by the average number of overlapping feedforward projections to any pair of neurons in the recurrent network (multiplied by their connection strength and *r*_*x*_), which gives (in block form)

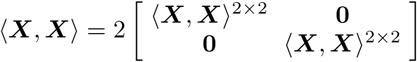

where **0** is the 2×2 zero matrix and **⟨*X, X* ⟩**^2^×^2^ is the external input CSD from the 3-population network, given by Eq. (8). Therefore, by Eq. (26),

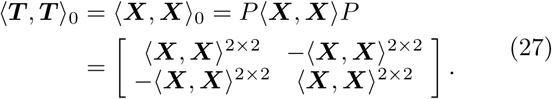

In other words, the mean total input CSD between excitatory neurons in the same subgroup (two neurons in *e*_1_ or two neurons in *e*_2_; diagonal blocks above) is positive and equal to half the mean external input between the same neurons. Hence, the cancellation by the recurrent network only reduces the external input CSD by a factor of 1/2, as opposed to the 𝒪𝒪 (1/*N)* reduction realized in the asynchronous state (when *W* is invertible). In contrast, the mean total input CSD between excitatory neurons in opposite subgroups (one neuron in *e*_1_ and the other in *e*_2_; off-diagonal blocks above) has the same magnitude as for same-subgroup pairs, but is negative. This represents a competitive dynamic between the two groups since they inhibit one another (recurrent connections are net-inhibitory in balanced networks [28, 45]), but receive different feedforward input noise. Interestingly, the average CSD between all pairs of spike trains is still 𝒪𝒪 (1/*N)* in this example, but it is easy to design examples with singular *W* in which this is not true.

Simulating this network for varying values of *N* shows that firing rates approach those predicted by the balance equation (23) (Fig. 5B), confirming that balance is realized. Pairs of excitatory neurons in the same group (both neurons in *e*_1_ or both neurons in *e*_2_) receive positively correlated external input and recurrent input (Fig. 5C, purple and green curves) that are partially canceled by negative correlations between their recurrent and excitatory input (Fig. 5C, yellow curve). Because the cancellation is only partial, the correlation between the neurons’ total inputs is 𝒪𝒪 𝒪(1) (Fig. 5C, black curve) in contrast to the asynchronous state (compare to Fig. 1G,H where cancellation is perfect at large *N)*. The total input covariance agrees well with the theoretical prediction from Eq. (27) (Fig. 5C, dashed gray line). As a result of this lack of cancellation between total input covariance, spike count covariances are also 𝒪 𝒪(1) and positive between same-group pairs (Fig. 5D). For opposite group pairs (one neuron in *e*_1_ and the other in *e*_2_), cancellation is also imperfect, but this leads to negative total input covariance, in agreement with the theoretical prediction from Eq. (27) (Fig. 5E), and leads to negative spike count covariances between neurons in opposite populations (Fig. 5F).

In summary, we have analyzed two mechanisms to generate 𝒪(1) spike train correlations in balanced networks. For the first mechanism (Fig. 2), spike trains in the external population are correlated so that external input correlations are 𝒪 𝒪(N). Cancellation is achieved so that spike train correlations are reduced to 𝒪(1). For the other mechanism (Fig. 5), external input correlation is 𝒪(1), but precise cancellation cannot be achieved so that spike trains inherit the 𝒪(1) correlations from the input. How could these two mechanisms be distinguished in cortical recordings? Under the first mechanism, we showed that fluctuations of inhibitory input to individual neurons closely tracks fluctuations of other neurons’ excitatory inputs (Fig. 3C). This should not be the case under the second mechanism because precise cancellation is not realized. Indeed, plotting the excitatory and inhibitory input to three excitatory neurons (two in *e*_1_ and one in *e*_2_) shows that input fluctuations are not closely tracked (Fig. 6). This provides a way to distinguish the two mechanisms from paired intracellular recordings. Indeed, the first mechanism (which we refer to as the “correlated state”) appears more consistent with the cortical recordings considered here (compare Fig. 3A to Figs. 3C and 6).

**FIG. 6.**
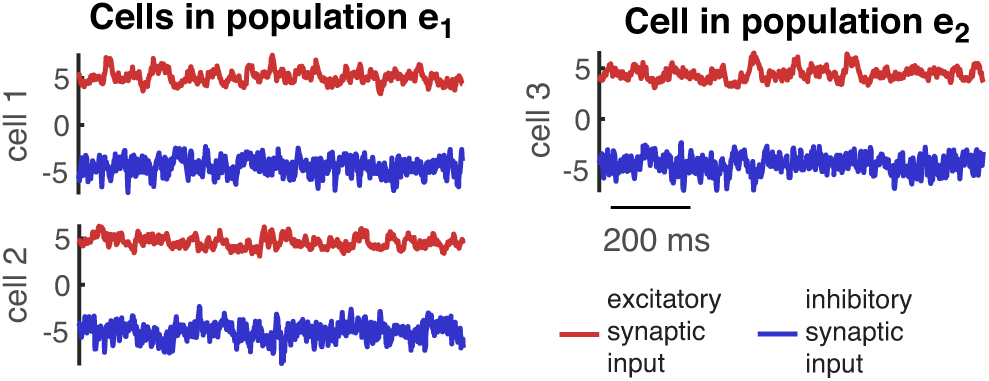
Synaptic input currents in a balanced network with correlations from singular mean-field connectivity. Same as Fig. 3A except for the network from Fig. 5. The left two traces are input currents to two excitatory neurons in population *e*_1_ (cells 1 and 2). The right traces are input currents to an excitatory neuron in population *e*_2_ (cell 3).

## VII. SUMMARY AND DISCUSSION

We analyzed correlated variability in recurrent, balanced networks of integrate-and-fire neurons receiving correlated feedforward input from an external population. We showed that correlations between spike trains in the recurrent network are small (𝒪 (1/*N)*) when spike trains in the external population are uncorrelated, consistent with previous work on the asynchronous state [31, 40], but much larger (𝒪(1)) when spike trains in the external population are correlated, giving rise to a “correlated state.” In both state⟨ *S, S* ⟩trong correlations in the feedforward input are canceled by recurrent synaptic input due to the excitatory-inhibitory tracking that arises naturally in densely connected balanced networks. In the correlated state, this cancellation allows for the derivation of a concise and accurate closed form expression for mean-field low frequency spike train CSDs and spike count covariances in terms of synaptic parameters alone. Hence spike count covariances in the correlated state are determined predominately by synaptic connectivity structure, not neuronal dynamics. The tracking of excitatory synaptic input by inhibition was observable on a pair-by-pair basis in the correlated state, but not the asynchronous state, suggesting that the correlated state is more consistent with some *in vivo* recordings.

In our analysis of the correlated state (*c >* 0), we only considered recurrent networks with two, statistically homogeneous neural populations: one excitatory and one inhibitory (with the exception of the simple multipopulation model analyzed in Section VI). Our analysis can be extended to arbitrarily many subpopulations as long as each sub-population contains 𝒪 𝒪(N) neurons, and also extends to networks with connection probabilities that depend on distance, orientation tuning, or other continuous quantities. This analysis has been developed for the asynchronous state in previous work [40, 41] and is easily extended to the correlated state as well. The primary difference is that **⟨*X, X* ⟩** is 𝒪 𝒪(N) instead of 𝒪(1).

Previous work has shown that networks with multiple sub-populations and networks with distance-dependent connectivity can break the asynchronous state in balanced networks when the network connectivity structure is constructed in such a way that the recurrent network cannot achieve the cancellation required for the asynchronous state [39–41], leading to 𝒪(1) correlations between some cell pairs. We showed that the precise tracking of excitation by inhibition provides an experimentally testable prediction for distinguishing this mechanism from the one underlying the correlated state (see Section VI).

Another alternative mechanism for achieving larger correlations in balanced networks is through instabilities of the balanced state. Such instabilities can create pattern-forming dynamics that produce intrinsically generated spike train correlations [42, 43, 62–68]. Some recordings show that local circuit connectivity can increase correlations [69], which is consistent with internally generated correlations, but inconsistent with the mechanisms that we consider here. In cultured populations of cortical neurons, moderate spike train correlations and excitatory-inhibitory tracking emerge even in the absence of correlated external input [30], suggesting that they are generated intrinsically. Correlations from pattern forming instabilities can be distinguished from the externally produced correlations considered here in at least two ways. Since instabilities generate correlations internally, they should produce weak correlations between activity in the recurrent network and activity in the external population(s) providing input to that network [42], in contrast to the mechanisms we consider here. Also, the presence of pattern forming instabilities in balanced networks depends on the timescale and spatial extent of excitatory and inhibitory recurrent synaptic projections as well as the strength of external input to the inhibitory population [42, 43, 62, 66, 68]. In cultured populations [30], these parameters can more easily be measured or modified to help determine the role of pattern forming instabilities in generating correlations.

We considered correlations produced by near-synchronous, correlated spiking in an external population (see Appendix C), but the correlated state could also be generated by time-varying firing rates in the external population. This could be modeled, for example, by replacing the homogeneous Poisson processes in the external population by doubly stochastic Poisson processed in which the rate fluctuations were shared across cells. This would produce 𝒪(N) external input covariance to the recurrent network and 𝒪(1) spike train correlation, so the mathematical analysis we considered applies equally well to this model. Future work should consider ways to distinguish correlations arising through shared rate fluctuations from those generated by near-synchronous spiking in a presynaptic layer.

In the correlated state, we found that the mean and empirical standard deviation of correlations is similar in magnitude, but in some recordings, mean correlations are much smaller than standard deviations [32, 35, 48, 70] suggesting that those circuits were in an asynchronous state, not a correlated state, during recording [48]. More generally, some recordings show very small mean correlations consistent with the asynchronous state [32, 48, 70], while others show larger correlation magnitudes [6, 33, 71]. Indeed, the magnitude of mean correlations can depend on many factors [6, 71], including arousal [72], attention [42, 73], stimulus [74], anesthetic state [31, 32, 35, 38], and cortical area and layer [34, 75–78].

In summary, spike train correlations and excitatory-inhibitory tracking likely arise from multiple mechanisms in different cortical areas, cortical layers, and conditions. The external input correlations introduced in this work are one such mechanism. Future studies should work to enumerate these mechanisms and generate experimentally testable predictions that distinguish them.

In the correlated state, spike train correlations in the recurrent network are essentially inherited from correlations between spike trains in the external population. Hence, the 𝒪(1) correlations realized by this mechanism require the presence of another local network with 𝒪(1) correlations. This raises the question of where the 𝒪(1) correlations are originally generated. One possibility is that they could be generated in a presynaptic cortical area or layer through the alternative mechanisms discussed above. Another possibility is that they originate from a network that is not in the balanced state at all. Nonbalanced networks can easily achieve 𝒪(1) spike train correlations simply from overlapping synaptic projections. While cortical circuits are commonly believed to operate in a balanced state, correlations could originate in thalamus, retina, or other sub-cortical neural populations then propagate to cortex.

The cancellation between variances of covariances observed empirically in simulations (Fig. 4F,H) is, to our knowledge, a novel observation, but we were unable to prove it analytically. Path integral approaches have recently been applied to compute variances of covariances in recurrent network models with uncorrelated external input [48]. Future work should consider the possibility of extending their analysis to networks with correlated external input.

Many previous studies of correlated variability in recurrent networks rely on linear response theory. In contrast, our derivation of Eqs. (9) and (19) used a weaker assumption that neural transfer of covariance is 𝒪(1) (Eq. (6) and surrounding discussion). Despite the fact that the derivation of Eqs. (9) and (19) do not depend on a linear response assumption, the equations themselves are linear and are the same that one would arrive at using linear response theory. Indeed, Eq. (9) and similar equations have been previously derived for various models using linear response techniques [5, 14, 31, 41, 46–48]. Hence, despite the use of neurons’ gains or susceptibility functions in this previous work, the resulting equations for mean-field covariance in the asynchronous state does not depend on the gains in the large *N* limit (see equations 38-39 in the supplement to [31] and equation 38 in [46]).

Even though the resulting equation is the same, our derivation of Eq. (9) is more general than derivations that rely on linear response theory. Specifically, linear response theory assumes that ⟨*S*_*a*_, *U*_*b*_⟩ *≈ A*_*a*_ ⟨*T*_*a*_, *U*_*b*_ ⟩, but our derivation is valid when relationship between *S*_*a*_, *U*_*b*_ and *T*_*a*_, *U*_*b*_ is nonlinear, as long as ⟨*S*_*a*_, *U*_*b*_⟩ / ⟨*T*_*a*_, *U*_*b*_⟩ ∼ *→* 𝒪(1) as *N* ∞ Our derivation also applies when ⟨*S*_*a*_, *U*_*b*_ ⟩/ ⟨*T*_*a*_, *U*_*b*_⟩ depends on the identity of *U, e.g*., it is different for *U* = *S* versus *U* = *X*. Our derivation is also valid when non-linear neural transfer causes crossfrequency coupling, *i.e*., when ⟨*S*_*a*_, *U*_*b*_⟩ (*f*_0_) depends on ⟨*T*_*a*_, *U*_*b*_⟩ (*f)* for values of *f ≠f*_0_. This is an interesting conclusion because frequencies are de-coupled in Eqs. (9) and, indicating an asymptotic linearity of the relationship between mean-field external input covariance and meanfield spike count covariance even in a non-linear system. This is possible because synapses in our model are linear and synaptic filtering dominates the network response properties at sufficiently large *N* and low frequencies (see Eqs. (6), (12), and surrounding discussion).

When input correlations are weak, covariance transfer for integrate-and-fire neurons is approximately linear so linear response approaches are justified [8, 53, 54], but transfer can become nonlinear (but still *O*𝒪(1)) when cor-relations are stronger [54, 79, 80]. Our results show that, even when the transfer of input covariance to spike train covariance is nonlinear, the mean-field relationship between **⟨*X, X* ⟩** and **⟨ *S, S* ⟩** is still linear to highest order in *N* and is the same relationship one would obtain if transfer were linear. Note that this linear relationship only applies at the mean-field level. Individual, pairwise CSDs between spike trains might still be non-linearly related to their pairwise external input CSDs. Recent work has called for looking beyond linear analysis of neuronal networks [81]. Our analysis shows that, even in networks where neural transfer of inputs is nonlinear, linear mean-field analysis can still be accurate and useful.

Despite the fact that Eq. (9) does not depend on neurons’ susceptibility functions or gains, it does depend on neurons’ mean spike count variance or power spectral densities. In contrast, Eq. (19) shows that mean-field covariance in the correlated state does not depend (to highest order in *N)* on neurons’ susceptibility functions, gains, power spectral densities, or spike count variance, but only involves synaptic parameters. This is similar to previous work showing that the dominant frequencies of oscillatory activity in balanced networks depends only on synaptic parameters [82]. Similarly, previous work showed that correlations in binary networks of inhibitory neurons without external drive are independent of connection strengths in the limit of strong connectivity (see equation 30 in [83] and equation 32 in [46]). The independence of spike count covariance on neurons’ susceptibilities and power spectral densities is an important conclusion because, for spiking neuron models, power spectral densities and susceptibility functions (or spike count variances and gains) can depend nonlinearly on the parameters of the neuron model as well as first, second, and higher moments of the neurons’ input statistics [84–87] and they can be difficult to compute numerically.

Numerical methods for approximating power spectral densities and susceptibility functions in networks of integrate-and-fire neurons have been developed using a diffusion approximation and Fokker-Planck techniques [8, 9, 11, 12, 49, 88], but this approach assumes that neurons’ synaptic input is dominated by or well-approximated by Gaussian white noise. Previous work satisfies this assumption by including Gaussian white noise as external input and making input from the recurrent network much weaker than this white noise input (for example, by making recurrent connectivity weak or sparse). In addition, instantaneous synapses (*η*_*b*_(*t*) = *δ*(*t*)) can make the diffusion approximation more accurate (although see [89, 90]). The diffusion approximation is not valid in our model since our external input is not well-approximated by Gaussian white noise, our recurrent input is approximately as strong as external input, and our synapses are not instantaneous. While some specialized approximations have been developed to avoid the assumption of Gaussian white noise input in various models and parameter regimes [1, 15, 55, 90–93], these approaches are outside the scope of our study since our central conclusion is that these approximations are unnecessary for deriving mean-field spike count covariance in the correlated state.

Three unproven assumptions underly our mean-field analysis of the correlated state. The first assumption is that neural transfer is 𝒪(1) (Eq. (6) and surrounding discussion). The third assumption is that mean-field power spectral densities or spike count variances {*S*_*a*_, *S*_*a*_} are 𝒪(1). The third assumption is that variability connection strengths is not strongly correlated with variability in individual CSD values (see comment in derivation of Claim 2 in Appendix A). These assumptions are made in other work, even if not stated explicitly. We have been unable to rigorously prove these assumptions for the model studied here, leaving an open problem for future work.

In summary, we showed that correlations in balanced networks can be caused by feedforward input from a population of neurons with correlated spike trains, defining the “correlated state.” In this state mean low-frequency CSD or spike count covariance is determined by a simple, closed-form equation of known parameters, greatly simplifying the analysis of spike count correlations in such networks. The correlated state predicts a precise balance between the fluctuations in excitatory and inhibitory synaptic input to individual neuron pairs, consistent with some in vivo recordings [23].

## ACKNOWLEDGMENTS

We thank Michael S. Okun for helpful comments on a draft of the manuscript and for his contribution to collecting the data used in Fig. 3A. We thank the reviewers for helpful comments that greatly improved the manuscript. RR was supported by National Science Foundation (https://www.nsf.gov/) grants DMS1517828, DMS-1654268, and Neuronex DBI-1707400. IL was supported by Deutsche Forschungsgemeinschaft Sonderforschungsbereiche (www.dfg.de/sfb) grant number 1089, Israel Science Foundation (https://www.isf.org.il/) grant numbers 326/07 and 1539/17, and Minerva (http://www.minerva.mpg.de/weizmann/). The funders had no role in study design, data collection and analysis, decision to publish, or preparation of the manuscript.

## Appendix A: Mean-field analysis of correlated variability in the asynchronous and correlated states

Here, we give a detailed derivation of mean-field CSDs that applies to both the asynchronous and correlated states. The only differences between parameters in those states is that ⟨*S*_*x*_, *S*_*x*_⟩ = 0 in the asynchronous state and ⟨*S*_*x*_, *S*_*x*_⟩ *#x223C;* 𝒪(1) in the correlated state. We derive CSDs in terms of ⟨*S*_*x*_, *S*_*x*_⟩ so that the results can be applied to either state. To more clearly organize the computation, we organize the calculation into the derivation of several claims that are derived separately. We start with a derivation of the mean-field external input CSD.

### Claim 1

*Mean-field external input CSD is given by*

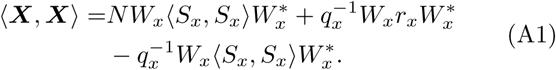

*Proof*. We first compute the external input CSD between neuron *j* in population *a* = *e, i* and neuron *k* in population *b* = *e, i* as

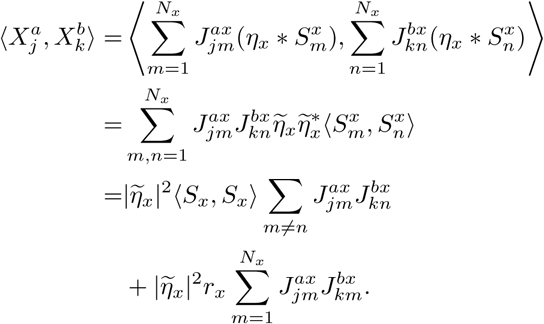

where we used the bilinearity of the Hermitian crossspectral operator, the fact that 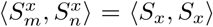 for *m ≠ n*, and 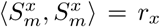 since external spike trains are Poisson processes. Averaging over *j* and *k* and using Eq. (2), we get

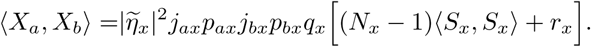

Writing this in matrix form and using 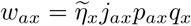 gives Eq. (A1).

Writing ⟨*S*_*x*_, *S*_*x*_⟩ ∼ *O*(*c*) so that *c* = 0 in the asynchronous state and *c* ∼ 𝒪(1) in the correlated state, Eq. (A1) implies that

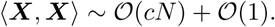

This expression helps understand the dependence of correlations on both *c* and *N*.

A similar derivation gives the mean-field CSD between neurons’ total and external inputs:

### Claim 2

*The mean-field CSD between external and total external inputs is given by*

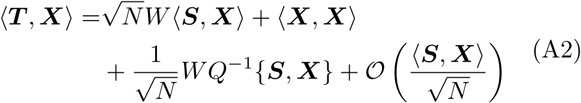

*Where*

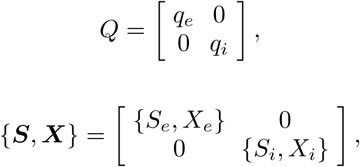

*and*

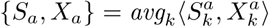

*quantifies the average CSD between neurons’ spike trains and their own external inputs*.

*Proof*. First break the total input into its constituent parts ***T*** = ***R*** +***X*** = ***E*** + ***I*** +***X***, so

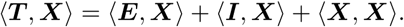

To compute the ⟨***E***, ***X* ⟩** term, we begin with

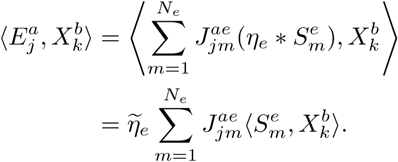

Before taking averages, note that whenever *b* = *e*, we need to treat the *m* = *k* term separately since it corresponds to the CSD between the spike train of and external input to the same neuron, which could scale differently than the case when *b* = *i* or *m ≠k*. Put another way, note that the definition of the mean-field values, ⟨*E*_*a*_, *X*_*b*_ ⟩ and ⟨*S*_*e*_, *X*_*b*_ ⟩, assumes that *m ≠ k* or *b ≠ e* (see Eq. (4)). Hence, computing averages separately for each of *b* = *e, i* gives

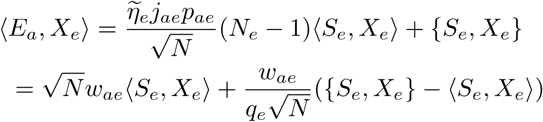

and

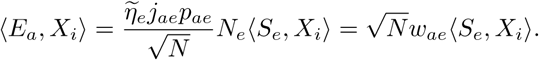

Note that this step requires us to assume that individual values of the random variable, 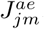, are not strongly correlated with individual values of 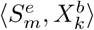, so that the expectation of their product can be replaced by the product of their expectations. This and similar assumptions are implicit in derivations in other studies, but are not typically proven or even explicitly stated.

An identical calculation for ⟨*I*_*a*_, *X*_*b*_ ⟩ gives

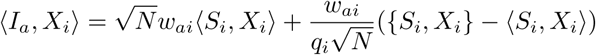

and

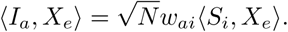

This allows us to write the CSD between recurrent and total input in matrix form as

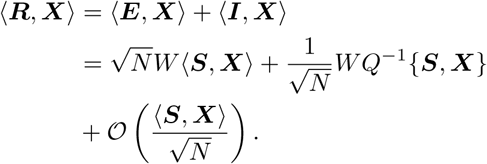

Now putting together ⟨***T, X* ⟩**= ⟨***R***, ***X* ⟩** + **⟨*X, X* ⟩** gives Eq. (A2).

A similar derivation gives the mean-field CSD between neurons’ total and external inputs:

### Claim 3

*The mean-field CSD between spike trains and total external inputs is given by*

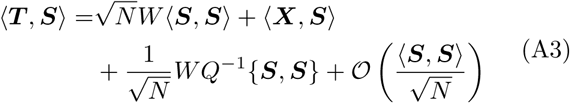

*where*

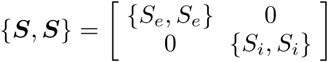

*and*

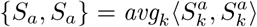

*quantifies the average power spectral densities of neurons’ spike trains*.

*Proof*. The derivation is identical to the derivation of Eq. (A2) above, but with *X* replaced by *S*.

We now use Claim 2 to derive the asymptotic behavior of ⟨***S***, ***X***⟩.

### Claim 4

*Under the assumption of O*𝒪(1) *transfer of meanfield covariance* (i.e., *Eq*. (7)*) and under the assumption that* {***S, S}*** ∼ *O*𝒪(1) (i.e., *neurons’ mean power spectral densities are O*𝒪(1)*), we have that*

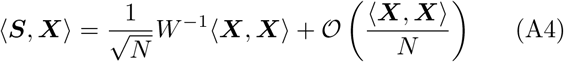

*in the limit of large N. For finite N, this approximation is only accurate whenever*

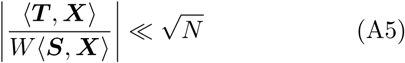

*where the division is performed element-wise*.

*Proof*. Combining the assumption of 𝒪(1) transfer of mean-field covariance (Eq. (6)) with Eq. (A2) gives

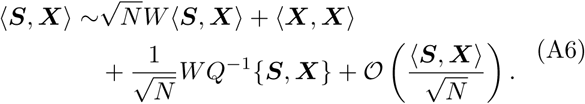

More rigorously, dividing both sides of Eq. (A2) elementwise by 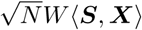 and applying Eq. (7) gives Eq. (A6). At finite *N*, note that this derivation of Eq. (A6) from Eq. (A2) is accurate whenever Eq. (A5) is true.

We do not know the exact scaling of {***S***, ***X*}** with *N*, but applying the Cauchy-Scwartz inequality gives

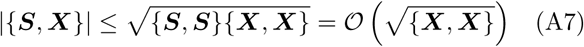

where we used our assumption that {***S, S}* ∼** 𝒪(1). To compute **{*X, X}***, we can repeat the derivation in Claim 1 with *j* = *k* to get

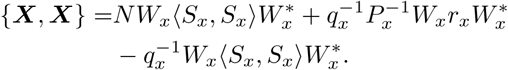

where

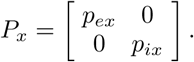

Comparing this to Eq. (A1), we may conclude that

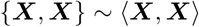

and therefore (using Eq. (A7) and the fact that **{*X, X* ⟩} *≥*** 𝒪(1)in both the asynchronous and correlated states), we have that

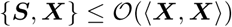

This allows us to rewrite Eq. (A6) as

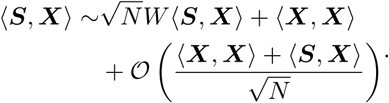

The only way that this equation can be self-consistent is if the terms on the right hand side cancel as *N* → ∞ which implies that 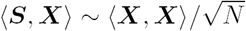 and, more specifically, that Eq. (A4) is satisfied as *N* → ∞.

We now derive the asymptotic values of mean-field spike train CSDs.

### Claim 5

*Under the assumptions made in Claim 4, we have that*

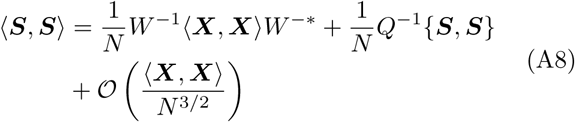

*in the limit of large N. For finite N, this approximation is accurate under Eq*. (A5) *and*

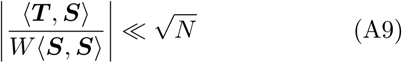

*where the division is performed element-wise*.

*Proof*. The proof is similar to that of Claim 4. Combining the assumption of 𝒪(1) transfer of mean-field covariance (Eq. (6)) with Eq. (A3) gives

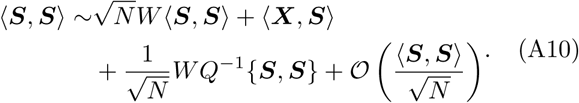

More rigorously, dividing both sides of Eq. (A3) elementwise by 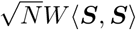 and applying Eq. (7) gives Eq. (A10). At finite *N*, note that this derivation of Eq. (A10) from Eq. (A3) is accurate whenever Eq. (A9) is true for ***U*** = ***S***.

Taking the conjugate-transpose of Eq. (A4) gives

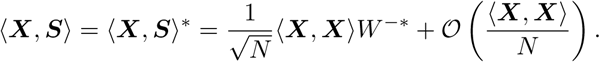

Plugging this into Eq. (A10) gives

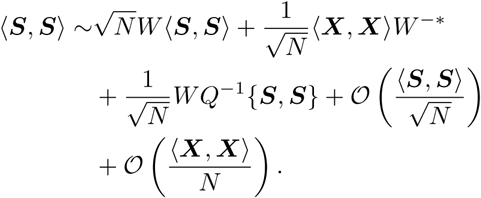

The only way this equation is self-consistent is if the terms on the right hand side cancel as *N* → ∞, which implies that **⟨ *S, S* ⟩ *𝒪*** (**⟨*X, X* ⟩**/*N* + **{*S, S}***/*N)* and, more specifically, Eq. (A8) is satisfied as *N*. For accuracy at finite *N* → ∞, we explicitly needed to assume Eq. (A9), but also implicitly assumed Eq. (A5) when we used Eq. (A4).

## Appendix B: Results from simulations with different connectivity parameters

So far, all of our simulations used the same connectivity parameters. Here, we consider results with different parameters. Specifically, we set *j*_*ee*_/*C*_*m*_ =35mV, *j*_*ei*_/*C*_*m*_ = *-*200mV, *j*_*ie*_/*C*_*m*_ = 120mV, *j*_*ii*_/*C*_*m*_ =*-*300mV, *j*_*ex*_/*C*_*m*_ = 200<u>m</u>V, and *j*_*ix*_*C*_*m*_ = 150mV. Note that *j*_*ab*_ was scaled by 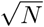 to produce the true connection strengths, as indicated in Results. We then repeated the simulations from Figs. 1I, 1J, and 2I. The result⟨ *S, S* ⟩hown in Fig. 7A,B, and C respectively, show similar overall findings to those reported for the connectivity parameters used in Results.

**FIG. 7.**
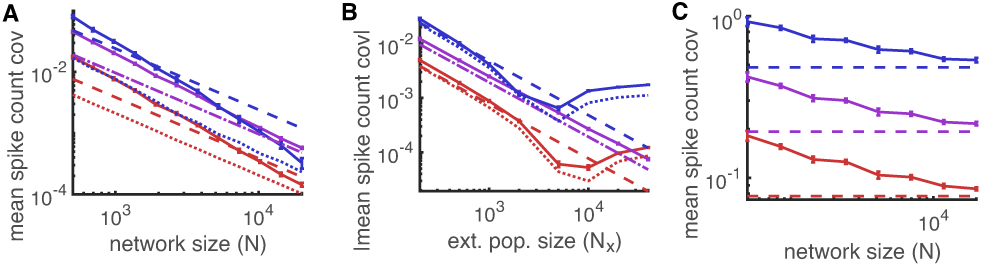
Results from simulations with different connectivity parameters. **A-C)** Same as Figs. 1I, 1J, and 2I respectively except that connectivity parameters were changed to *jee*/*Cm* = 35mV, *jei*/*Cm* = *-*200mV, *jie*/*Cm* = 120mV, *jii*/*Cm* = *-*300mV, *jex*/*Cm* = 200mV, and *jixCm* = 150mV.

## Appendix C: Generation of correlated spike trains for external inputs

To generate correlated, Poisson spike trains for the external population in the correlated state we used the multiple interaction process (MIP) method [84] with jittering. Specifically, we generated one shared “mother” process with firing rate *r*_*m*_ = *r*_*x*_/*c*. Then, for each of the *N*_*x*_ “daughter” processes, we randomly kept each spike in the mother process with probability *c*. As a result, each daughter process is a Poisson process with firing rate *cr*_*m*_ = *r*_*x*_ and a proportion of *c* of the spikes are shared between any two daughter processes. To get rid of perfect synchrony between the daughter processes, we jittered each spike time in each daughter process by a normally distributed random variable with mean zero and standard deviation *τ*_*c*_ = 5ms. Upon jittering, the daughter processes remain Poisson and the resulting CSD between daughter processes is given by Eq. (20). Spike count correlations between the daughter processes over large time windows are exactly *c*. The daughter processes were used as the spike trains, 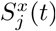 in the external population in the correlated state. See [84] for a deeper analysis of this algorithm.

## Appendix D: Parameter values and details of computer simulations

All connection probabilities were *p*_*ab*_ = 0.1 for *a* = *e, i* and *b* = *e, i, x*. Synaptic timescales were *τ*_*e*_ = 8ms, *τ*_*i*_ = 4ms, and *τ*_*x*_ = 10ms. The firing rate of the external population was *r*_*x*_ = 10Hz and, in the correlated state, the correlation was *c* = 0.1 with a jitter of *τ*_*c*_ = 5ms. All covariances and correlations were computed by counting spikes or integrating continuous processes over a window of length 250ms. Membrane capacitance, *C*_*m*_, is arbitrary so we report all current-based parameters in relation to *C*_*m*_. For convenience, one can therefore set *C*_*m*_ = 1. Unscaled connection strengths were *j*_*ee*_/*C*_*m*_ = 25mV, *j*_*ei*_/*C*_*m*_ = *-*150mV, *j*_*ie*_/*C*_*m*_ = 112.5mV, *j*_*ii*_/*C*_*m*_ = *-*250mV, *j*_*ex*_/*C*_*m*_ = 180m<u>V</u>, and *j*_*ix*_*C*_*m*_ = 135mV. Note that *j*_*ab*_ was scaled by 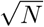 to produce the true connection strengths, as indicated in Results. Neuron parameters are *g*_*L*_ = *C*_*m*_/15, *E*_*L*_ = −72mV, *V*_*th*_ =-50mV, *V*_*re*_ = −75mV, *V*_*lb*_ = − 100mV, Δ_*T*_ = 1mV, and *V*_*T*_ = − 55mV. Synaptic currents in figures are reported in units *C*_*m*_*V*/*s*. Covariances between synaptic currents are computed between integrals of the currents (see Eq. (3) and surrounding discussion), so the covariances have units 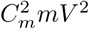.

The differential equations defining the neuron model were solved using a forward Euler method with time step 0.1ms. Statistics in Figs. 1D, 2C,D, and 4E,F were computed from a simulation of duration 50s. Statistics in Figs. 1E-I, 2E–I, and 4C,D,G,H were computed by repeating a simulation of duration 50s over ten trials for each value of *N*, then averaging over trials. For each trial, network connectivity was generated with a different random seed, so the statistics are averaged over time and over realizations of the “quenched” variability of network connectivity.

